# Sex differences in behaviour and molecular pathology in the 5XFAD model

**DOI:** 10.1101/2021.04.29.440396

**Authors:** Annesha Sil, Arina Erfani, Nicola Lamb, Rachel Copland, Gernot Riedel, Bettina Platt

## Abstract

**Background:** The prevalence of Alzheimer’s Disease (AD) is greater in women compared to men, but the reasons for this remain unknown. This sex difference has been widely neglected in experimental studies using transgenic mouse models of AD.

**Objective:** Here, we studied behaviour and molecular pathology of 5-month-old 5XFAD mice, which express mutated human amyloid precursor protein and presenilin-1 on a C57BL/6J background, vs. their wild-type littermate controls, to compared both sex- and genotype-dependent differences.

**Methods:** A novel behavioural paradigm was utilised (OF-NO-SI), comprising activity measures (Open Field, OF) arena, followed by Novel Object exploration (NO) and Social Interaction (SI) of a sex-matched conspecific. Each segment consisted of two repeated trials to assess between-trial habituation. Subsequently, brain pathology (amyloid load, stress response and inflammation markers, synaptic integrity, trophic support) was assessed using qPCR and Western blotting.

**Results:** Female 5XFAD mice had higher levels of human APP and beta-amyloid (Aβ) and heightened inflammation vs males. These markers correlated with hyperactivity observed in both sexes, yet only female 5XFAD mice presented with deficits in object and social exploration. Male animals had higher expression of stress markers and neurotrophic factors irrespective of genotype, this correlated with cognitive performance.

**Conclusion:** The impact of sex on AD-relevant phenotypes is in line with human data and emphasises the necessity of appropriate study design and reporting. Differential molecular profiles observed in male vs. female mice offer insights into possible protective mechanisms, and hence treatment strategies.

## INTRODUCTION

As the most common cause of dementia, Alzheimer’s Disease (AD) is a progressive debilitating neurodegenerative disease currently affecting > 500,000 people in the UK. Dementia is characterised by progressive cognitive deficits, memory loss and non-cognitive behavioural changes in, for example, activity, sleep, and mood [1].

The incidence for AD appears to be greater in women compared to men even when corrected for demographic differences [2, 3]; AD-associated hippocampal atrophy and cognitive decline are also more prominent in female patients [4]. This was traditionally attributed to the longer lifespan of women, but recent research has indicated that biological differences (for example in chromosomes, hormones or epigenetics) may be just as important [5–7]. Even though human studies have provided some insight into these sex differences in AD patients, sex differences in experimental models of AD continue to produce varying and sometimes conflicting results, especially when trying to associate behavioural and molecular pathologies. For example, female 3xTg-AD mice (with transgenes for APP (amyloid precursor protein, Swedish mutation), tau (P301L mutation), and PSEN-1 (presenilin-1, M146V mutation) displayed greater spatial cognitive deficits, neuroinflammation and Aβ burden relative to males of the same age [8, 9]. On the other hand, sexual dimorphism noted in P301S Tau transgenic animals indicated that male mice displayed significant changes in a composite behavioural phenotype as well as in tau phosphorylation. Females presented with impairments only in specific behavioural tests, such as Morris water maze and open field and had lower expression of some inflammatory markers [10]. Conversely, no sex differences in gene expression were observed in other models such as our knock-in PLB1_triple_ mice (with transgenes for APP, tau and PSEN1) or the PLB4 (knock-in of human BACE1 (β-secretase)) mice [11, 12].

Neuropathologically, AD is characterised by the cleavage and subsequent processing of two proteins, i.e. APP and the microtubule-associated protein tau [13]. APP is preferentially cleaved by BACE1, leading to the loss of secreted APPα in favour of beta-amyloid (Aβ) [14], which aggregates into oligomers and fibrils causing the formation of plaques. It is generally assumed that metabolites of APP (and tau) are toxic and cause neuroinflammation, abnormal stress of the endoplasmic reticulum (ER), and dysregulation of neurotrophic factors, jointly leading to neurodegeneration [15].

Chronic neuroinflammation is characterised by an increase in activated astrocytes and resident microglia, but also by the activation of the NLRP3 (NOD-, LRR- and pyrin domain-containing protein 3) inflammasome, and these appear to trace disease progression [16, 17]. Particularly relevant here are immunoreactive Iba-1 (Ionized calcium binding adaptor molecule 1) and GFAP (Glial Fibrillary Acidic Protein) as markers for activated microglia and astrocytes respectively, which occur in close proximity to Aβ deposits [18–20]. NLRP3, on the other hand, has shown promise as an experimental therapeutic target [21, 22]. Similarly, protein aggregation in AD may be associated with the chronic unfolded protein response, typical for prolonged stress resulting in apoptosis and synaptic loss [15, 23]. This pathway is connected to amyloid pathology since tunicamycin-induced ER stress can cause overproduction of Aβ peptides in cell lines [24] while conversely, Aβ is known to increase multiple ER stress markers [25]. Repair and protective mechanisms in neurones are in place to counteract these damaging events, such as brain-derived neurotrophic factor (BDNF), tropomyosin-related kinase B (TrkB), and cAMP response element-binding protein (CREB), and these are all compromised by Aβ accumulation [26, 27].

Overall, how cellular stress and inflammation relate to toxic proteins such as Aβ, and more specifically, whether interactions are affected by sex, still remains elusive. Here, we employed one of the most commonly used murine models of amyloid pathology, the 5X Familial AD mouse (5XFAD), which expresses the three human APP (hAPP) gene mutations alongside two human presenilin-1 (hPSEN1) mutations [28]. From the three lines originally generated, Tg7699 (now termed 5XFAD) expressed the highest levels of transgenic APP and displayed early (at 2 months) and aggressive amyloid pathology, as well as elevated inflammatory markers (GFAP) and reduced levels of synaptic markers (synaptophysin and PSD-95) by 9 months of age [28]. Surprisingly, an investigation into ER stress markers in a mixed-sex cohort in this model failed to identify genotype differences [29]. However, conditional delivery of BDNF from astrocytes rescued memory impairments and prevented synaptic loss in 5XFAD mice [30].

The impact of sex on phenotypes in 5XFAD mice is still controversial. While Sadleir and co-workers [31] recorded heightened levels of Aβ42 in female transgenic animals relative to their age-matched male counterparts, a transcriptomic and proteomic analysis of the hippocampi of 4 month old 5XFAD animals did not reveal any sex biases for the development of amyloid pathology or the inflammation marker GFAP [32]. However, female 5XFAD mice showed heightened gliosis in both cortex and hippocampus aged 13 months relative to males [33] and sustained microglial activation from 4 to 9 months of age [34].

More conflicting are data on the behavioural profile of 5XFAD mice. Male transgenic mice presented with working and short-term memory deficits aged 4-5 months, but there was no overall change in locomotor activity [20, but see 28 for contrasting results]. However, deficits in activity were reported in specific motor tasks from 9 months, but this was not sex specific [35]. Ambulation in the open field or Y-maze was also reduced at this age in females only [36], but this phenotype was lost in older cohorts [33, 37]. Data from 12-month old male 5XFAD mice suggested intact memory during object recognition tasks in an open field and Y-maze while age-matched females displayed clear impairments [33]. The authors suggested that the study design and complexity of the task are determinants for the discrimination of sex-specific phenotypes. Studies into social deficits, while limited, have indicated abnormal social recognition at 9 months of age (mixed sex; [38] as well as decreased exploration of conspecifics in an age-dependent manner in females only [39]). Differences in test parameters as well as appropriate study design choices of controls [40] have likely contributed to the reported discrepancies. Finally, many studies do not report the sex of animals used [41]. And it is frequently unclear whether 5XFAD animals used were on a C57BL6/J background, and whether the recessive retinal degradation allele *Pde6b^rd1^* or the muscular dystrophy gene *Dysf^im^* found in the original SJL background were eliminated [35].

Here, we used our novel OF-NO-SI paradigm, which combines elements of the open field, novel object recognition and social interaction [42]. Its reliance on multiple neurotransmitters makes it an ideal candidate task to explore genotype- and sex-related phenotypes, and to confirm whether such phenotypes are consistent with previously reported data from 5XFAD mice. *In vivo* testing was followed by a detailed within-subject post-mortem tissue analysis to measure amyloid pathology, inflammatory, synaptic, ER stress and neurotrophic markers, and explore correlations between behavioural and molecular endpoints.

Our data suggest distinct behavioural and molecular phenotypes in female vs. male 5XFAD littermates, with only the former displaying overt behavioural anomalies that correlated with heightened levels of beta-amyloid and inflammation, while the latter showed an upregulation of potentially neuroprotective pathways.

## MATERIAL AND METHODS

This work was conducted according to the ARRIVE guidelines 2.0 for reporting animal research [43] and the recommendations of the EQIPD (European Quality in Preclinical Data) WP3 (Work Package 3) study group for the internal validity in the design, conduct and analysis of preclinical biomedical experiments involving laboratory animals [44].

### Animals and experimental conditions

5XFAD (B6.Cg-Tg(APPSwFlLon,PSEN1*M146L*L286V)6799Vas/Mmjax) mice on BL/6J background were purchased from The Jackson Laboratory (Bar Harbor, Maine, USA: stock number 34848-JAX) and bred and maintained on-site at the University of Aberdeen by backcrossing to C57BL6/J mice. Genotypes were confirmed from ear-biopsies to identify transgenic (5XFAD) animals expressing both APP and PSEN-1 and wildtype (WT) genes, carried out by Transnetyx Inc. (Cordova, USA). A total of 40 five-month-old mice (male WT, n=10, female WT, n=10; male 5XFAD, n=10; female 5XFAD, n=10) were utilised. Animals were group-housed together in conventional type 3 polycarbonate stock cages with a maximum of 7-8 animals per cage, separated by sex, and maintained on a 12-hour day-night cycle (lights on 7am, simulated dusk/dawn 30 mins) in temperature- (20-22°C) and humidity- (60-65%) controlled holding facilities with ad libitum access to both food (Special Diet Services, Witham, UK) and water. Enrichment was provided by paper strips, cardboard tubes, and corncob as bedding (DBM Scotland Ltd). Behavioural testing took place on weekdays during the light period in a separate testing room. Three age- and sex-matched C57BL/6J mice (either male or female) served as stranger mice for social interaction trials and were kept in a separate holding room, under the same housing conditions. The experimenter was blinded to the genotype of the subject and testing order was randomised using a random number generator (www.random.org). No power calculation was available; estimates were based on previous research [42]. All procedures were approved by local ethical review, a UK Home Office project licence and complied with the EU directive 63/2010E and the UK Animal (Scientific Procedures) Act 1986.

### Open Field-Novel Object-Social Interaction (OF-NO-SI) Paradigm

Experiments were conducted in a white circular Perspex arena of 50cm diameter, illuminated by white LED lights (average light intensity recorded in the arena =160.6 lux) facing directly downwards onto the arena. Animals were tracked using an overhead camera (Sony) connected to the ANY-maze video and activity tracking software (v 6.0, Ugo Basile, Comero, Italy) complemented by manual observations by the experimenter for urination and defecation during different trials but no differences were observed (data not reported) [42]. Between trials, the apparatus was cleaned with non-alcohol, fragrance-free wipes between trials to remove odour cues.

Prior to experimentation, mice were individually housed in transport cages (conventional shoe box cages (Tecniplast, Italy) with fresh bedding, water but no food) and habituated to the testing room for 45 minutes. Average temperature and humidity of the testing arena were recorded as 20.5°C and 60.5%, respectively. Arena definitions included a wall zone (5cm from the outer wall of the arena) and an interaction zone (centric 5cm wide ring around the cylinder (Fig. 1C)). The protocol followed exactly the one reported by Yeap et al. [42] and consisted of three test stages, each with 2 trials of 10 mins, separated by an inter-trial interval (ITI) of 15 minutes, during which the test animals were returned to their transport cages (Fig. 1A and 1B). The first stage consisted of two open field trials (OF); in the first (OF1) the animal was placed randomly along the wall of the empty arena and allowed to explore freely for 10 minutes. This was repeated for the second open field trial (OF2) after the 15-minute ITI. OF trials served as a habituation component to the remainder of the task components.

**Fig. 1:**
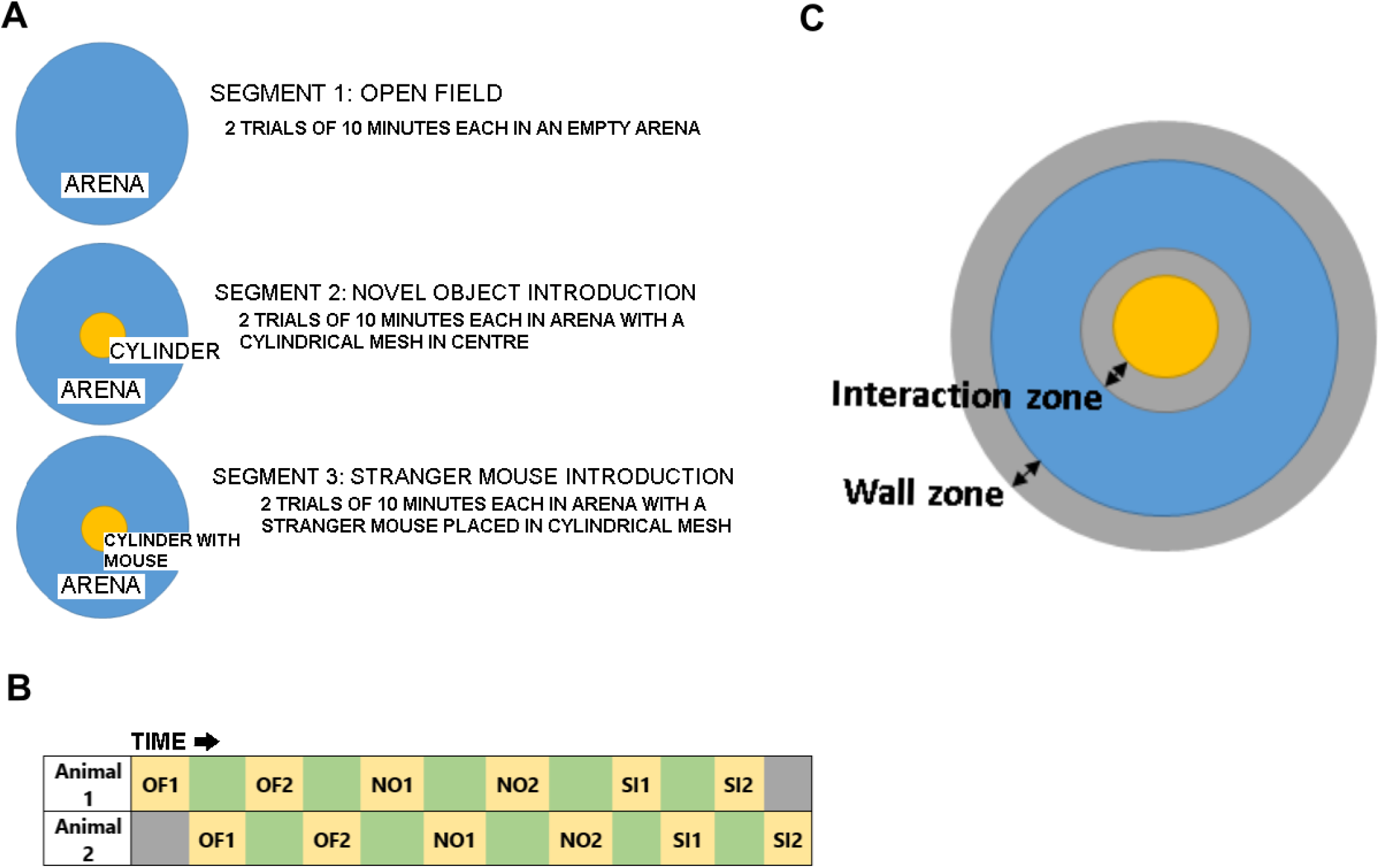
The OF-NO-SI experimental design: OF-NO-SI consists of a circular Perspex arena first used as an open field (OF) **(A)**. Next, a cylinder is placed in the centre of the arena (novel object trials, NO) and finally a stranger mouse was introduced into the cylinder for social interaction trials (SI). **(B)** An overview summary of a single run of the OF-NO-SI experiment: green boxes depict inter-trial intervals (ITIs, 15- minutes), during which a second animal can be interspersed. **(C)** An outline of borders and zones used for the parameters examined: The ‘wall zone’ (5cm from the outer wall) and ‘interaction zone’ (5cm from the cylinder) are indicated.

In the second stage, a novel object (cylindrical wire-mesh cage: 8 cm in diameter, 20 cm tall) was introduced into the centre of the arena. Again, two trials (NO1, NO2) were administered, all other parameters were identical to the OF trials. In the third social interaction stage, a stranger mouse was introduced into the cylinder; all other parameters were identical to OF and NO stages. Animals were run in pairs so that one animal was tested while the other had its ITI (Fig. 1B).

### Behavioural data analyses

The distribution of the animal in the arena was visualised by generating heat maps [42] using MatLab R2017a (MathWorks, Massachusetts, United States), which illustrated average location preference during each trial for one representative animal per genotype and sex. Primary behaviours (activity measured as distance moved, time spent in predefined zones and number of visits to selected zones) were recorded and analysed using ANY-maze behavioural tracking software (Ugo Basile, IT) tracking the centre of gravity at 10Hz sampling rate. One female (5XFAD, no 371) had to be excluded from analysis because of a tracking error.

Data are presented as parameters of interest averaged over the 10-minute duration of each trial. Total path length (in m) in 10 minutes of each trial during OF was recorded as a measure of locomotor activity while time (s) in wall zone served as a proxy for anxiety related behaviours. To establish effective habituation to the open field, data for OF trials are presented as distance moved per minute (see Supplementary Information). Time (in s) spent in the interaction zone (<5 cm from the cylindrical mesh cage; Fig. 1C) was recorded during NO and SI phases and represents exploration of the object/stranger. The number of visits to the interaction zone was also recorded as a proxy of purposeful approaches to the object / stranger.

Statistical analysis and graphs were prepared using Prism (V 8.0; GraphPad; USA). Data were averaged per genotype and sex and are represented as mean ±SD for each trial. Normality of data was confirmed using the Shapiro-Wilk normality test, while outliers were determined using the method of Grubbs (3- standard deviation from mean). This led to the exclusion of one female WT (no. 384) which showed abnormal hyperactivity during all phases of the paradigm. For all analyses, alpha was set to 5% with trends beings reported at p<0.09.

Main effects and interactions were assessed for each stage using a three-way analysis of variance (ANOVA) with sex (male or female), genotype (5XFAD vs WT) and trial (repeated measures, RM) as factors. *Within-trial* habituation (indicated by declining values over the 10 minutes during trial 1) and *between-trial habituation* (indicated by higher activity during trial 1 vs trial 2) was probed using a 2-way ANOVA for each group with trial and time as repeated measures for specifically the OF stage (see Supplementary information). Familiarity indices were also calculated for NO and SI stages [(Time in/Number of visits to interaction zone in trial 1) - (Time in/number of visits to interaction zone in trial 2)]/(Total time in/total visits to zone) to evaluate memory formed during trial 1; positive values constitute recall. For the familiarity indices during NO and SI, a one-sample t-test against a hypothetical chance value of 0 was employed within each genotype of each sex to observe if there was a significant change in either time spent in zone/number of visits between trial 1 and trial 2. Group effects were determined using 1-way ANOVA.

### Brain extraction and lysate preparation

Mice utilised in the OF-NO-SI experiment were terminally anaesthetised with Euthatal (Merial Animal Health Ltd., Lyon, France) intraperitoneally and transcardially perfused with 0.9% heparinised saline. Extracted hemi-brains were snap-frozen in liquid nitrogen and stored in −80°C until use. Samples were processed as described previously [45] to obtain soluble and insoluble fractions. In brief, brain tissue (∼80mg) was homogenised in ∼1:10 (w/v) Igepal (Sigma, Dorset, UK) based non-denaturing, non-ionic lysis buffer (in mM: 20 HEPES, 150 NaCl, 1% Igepal, 0.1 EDTA, pH= 7.6) with inhibitors for protease and phosphatase (PhosStop) (Roche Life Science, Burgess Hill, UK). Samples were centrifuged (13,000g, 4 °C for 20 mins); the supernatant constituted the soluble fraction. The pellet was homogenised twice in Igepal lysis buffer (1ml) via repeated aspiration with a 1ml pipette tip, briefly vortexed and then centrifuged (14,000 rpm, 20 mins), and then resuspended in 1:1 (w/v) 70% formic acid at 4 °C for overnight agitation. The formic acid fraction was then spun (18000g, 4 °C for 20 mins) and the resultant supernatant formed the insoluble fraction. Before use, the insoluble fraction was mixed with 4 volumes of neutralising buffer (2M Tris + 2M NaH2PO4).

### Immuno-blotting

Western and dot blots were performed as previously described [45]. Protein concentration for the soluble fraction was adjusted (3µg/ml) following bicinchoninic acid assay measurement (BCA, Sigma-Aldrich, Poole, UK) and dilution in lysis buffer. A set volume of neutralised samples for the insoluble fraction was utilised as protein concentration evaluation using BCA is not possible due to the strong reducing action of formic acid [46].

For Western blotting, all samples (soluble and insoluble) were mixed with lithium dodecyl sulphate (LDS, Thermo Fisher, Paisley, UK) and 15 mM dithiothreitol (DTT, Sigma), and heated for 10 mins at 70 °C (apart from samples meant for detection of amyloid) and run on 4–12% Bis-Tris precast gels (Nupage, Thermo Fisher Scientific) in MOPS buffer (Nupage, Thermo Fisher Scientific) for 45 mins (or MES buffer (Nupage Thermo Fisher Scientific) for 35 min in case of Iba-1 only) at 200V constant voltage. For detection of full-length APP (fAPP) and monomeric Aβ, samples were mixed only with LDS and loaded onto the precast gels in MES buffer without an additional heating step. For these amyloid markers and Iba-1, transferring was done onto 0.2 µm nitrocellulose membranes via the iBlot system (Nupage, Thermo Fisher Scientific), followed by microwaving for 3 mins in PBS (phosphate buffered saline (Sigma Aldrich, UK) for detection of low molecular weight proteins; for all other proteins, standard transfer conditions were applied onto 0.45 µm nitrocellulose membranes.

For the selective detection of amyloid-beta (Aβ) via dot blots [45], samples were dotted directly onto 0.2 µm nitrocellulose membranes (5 µl of 2 µg/µl per dot for the soluble fraction and 5 µl of neutralised insoluble fraction samples). No DTT, LDS or heating steps were involved. Following this, washing of all blots was carried out in 0.05% Tween-20 (Sigma) containing Tris-buffered saline (TBST; in mM: 50 Trizma base, 150 NaCl, pH= 7.6). All membranes were then blocked for 1 hour at room temperature in TBST containing 5% milk powder. Primary antibodies in 5% bovine serum albumin (BSA, Sigma Aldrich, UK) containing TBST were then added to the membranes for overnight incubation at 4 °C (refer to Table 1 for a list of antibodies utilised). Secondary antibodies (as appropriate) were added for 1 hour at room temperature (goat anti-rabbit/ goat anti-mouse, IgG, HRP conjugated; Merck Millipore (1:5000)) before visualisation using enhanced chemiluminescence (1.25 mM luminol, 30 µM coumaric acid, 0.015% H_2_O_2_). A Vilber-Fusion-SL camera (Vilber, Eberhardzell, Germany) and iBright™ FL1000 Imaging System camera (Invitrogen™, ThermoFisher Scientific) were utilised to capture immunoreactivity at 16bit for analysis. Membranes were then processed for total protein using both Ponceau and Coomassie protein stains as described previously [45].

**Table 1.**
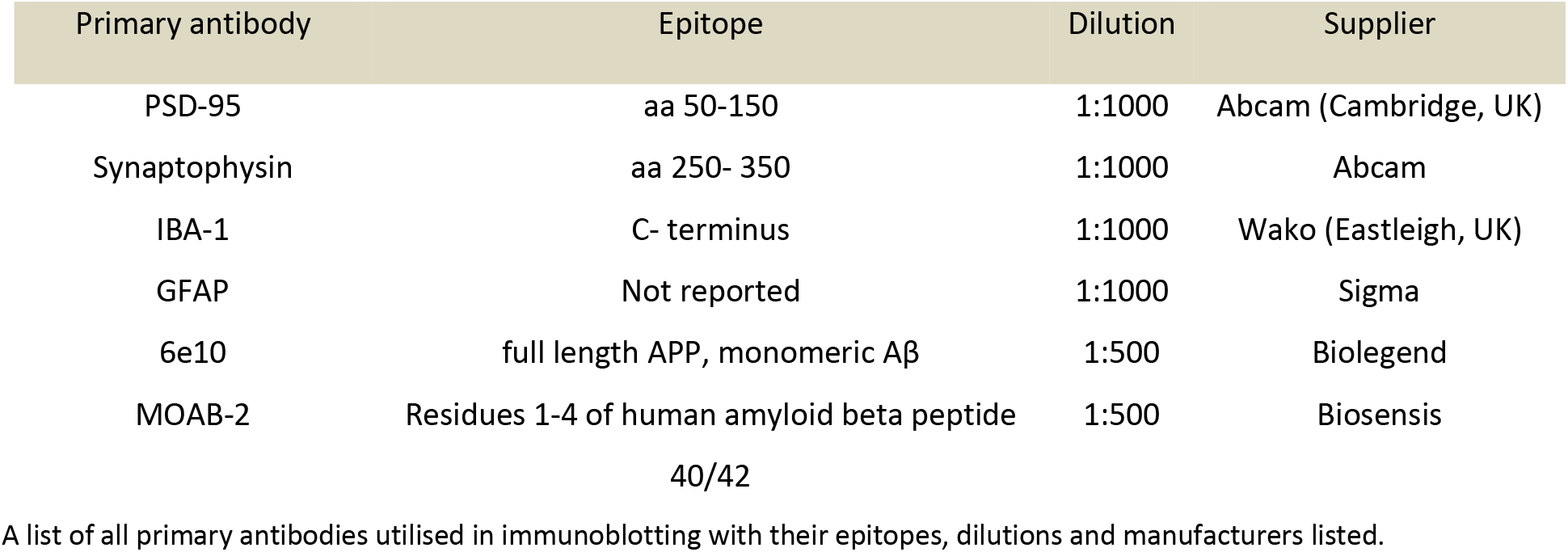
Primary Antibodies utilised for Western blotting.

### qPCR

Total RNA was extracted from snap-frozen hemi-brain tissue using TRI Reagent (Ambion, Warrington, UK) as per the protocol of the manufacturer. Bioline cDNA synthesis kit (Bioline, London, UK) was utilised to synthesize cDNA from 1 μg of total RNA. Amplification of genes of interest was achieved by quantitative polymerase chain reaction (PCR) in a Roche LightCycler® 480 System (Roche Diagnostics, Burgess Hill, UK) using GoTaq qPCR Master Mix (Promega, Southampton, UK). Relative gene expression was calculated using the comparative Ct method (2−δδCt)[47]. A comprehensive list of all primer sequences utilised can be found in Table 2. Normalisation of the data utilised the geometric mean of three of the most stable reference genes (Y-Whaz, NoNo, 18S, GAPDH or BetaActin). As human APP (hAPP) and PSEN-1 (hPSEN-1) were not detectable in WT animals, a ratio of the Ct values of most stable reference gene on the plate and the gene of interest was carried out to obtain gene expression data.

**Table 2.**
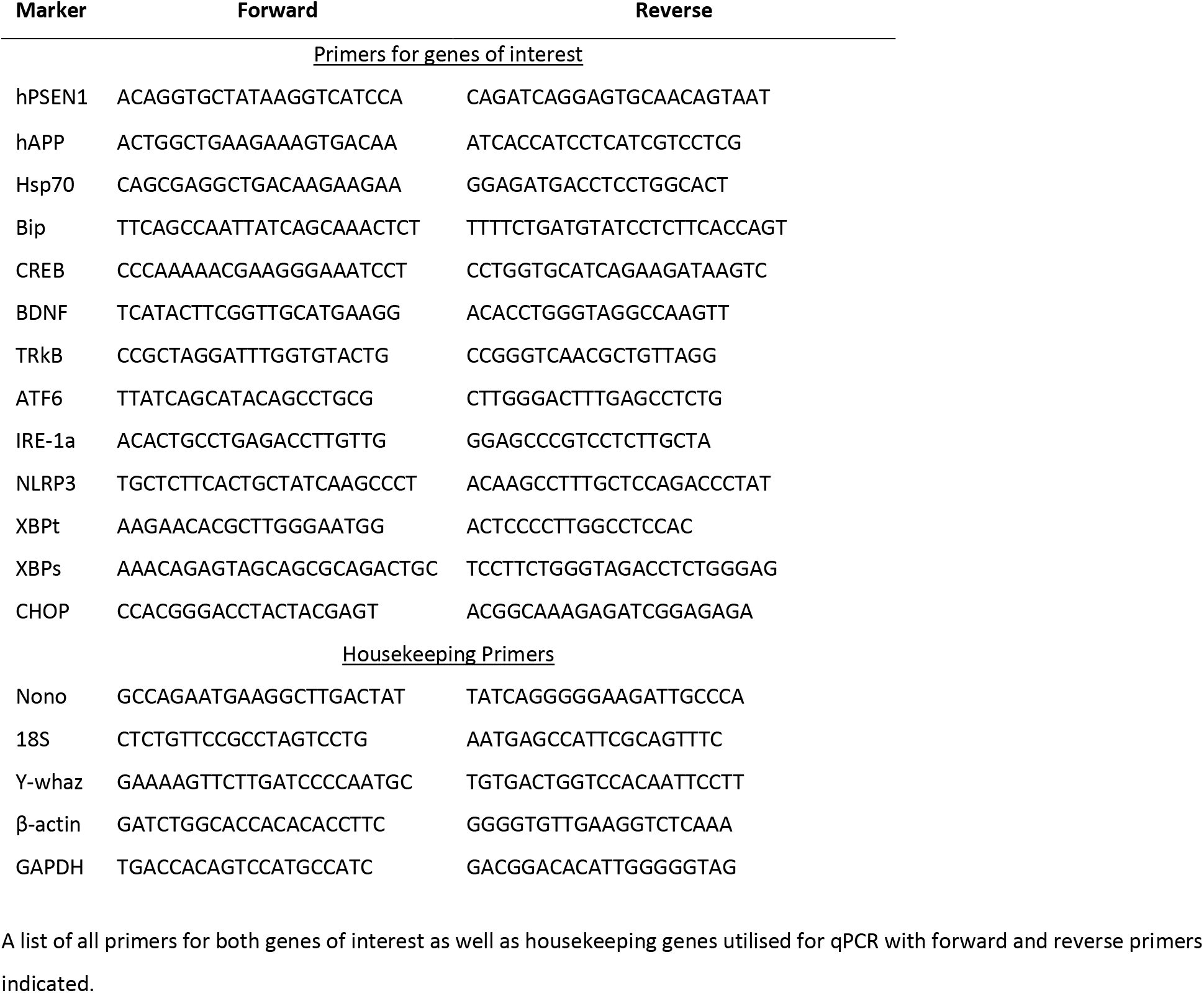
qPCR Primers.

### Analysis of molecular data

Quantification for Western and dot blots was performed as area under the curve (AUC) using ImageJ (Ver. 1.51, NIH, USA) software, and adjusted to total protein as per Coomassie or Ponceau. Fold change was then calculated relative to female WT controls (the WT control group was selected randomly). Statistical analysis was carried out using Prism (V8.0, GraphPad). Normality was assessed using a Shapiro-Wilk normality test and outliers were determined using the Grubb’s test. Unless otherwise stated, data sets were analysed using a regular two-way ANOVA to measure the effect of genotype and sex with selected post-hoc paired comparisons using Bonferroni correction. For gene expression data of hAPP and hPSEN1 in the transgenic animals, an unpaired parametric t-test was carried to observe differences between sexes. Data are presented as bar charts with scatter and alpha was set to 5%. Only significant terms (p<0.05) and trends (p<0.09) are given for clarity.

### Correlations

Correlational analysis was carried out between behavioural and molecular markers of interest. For the behavioural markers, total activity in OF, NO and SI, as well as time in wall zone during OF, time in zone and number of visits during NO and SI and NO and SI familiarity indices for time in zone (referred to as NO and SI index) were utilised. All markers underwent Z-transformation within each group (combined♀+♂ 5XFAD, ♀5XFAD, ♂5XFAD, ♀+♂ WT, ♀WT, ♂WT) followed by Pearson’s correlation analyses after confirmation of Gaussian distribution of datasets. Correlations are displayed in the form of a heat plot matrix with red indicating negative correlations and blue indicating positive ones. For all data, significance was set at p<0.05, with trends being reported as p<0.09.

## RESULTS

### OF-NO-SI Behavioural Experiments

#### Normal habituation amidst heightened activity in 5XFAD mice

Exploration of the open field was recorded as activity, defined as total distance moved, during the entire length of OF1 or OF2. A 3-way ANOVA with sex, genotype and trial (RM) as factors revealed a very strong effect of genotype (F (1,34) = 18.93); p<0.0001), due to higher locomotor activity in 5XFAD animals compared to WT (Fig. 2A). A strong effect of trial (F (1,34) = 182.0; p<0.0001) confirmed lower activity during OF2 compared to OF1, i.e. habituation between trials. And a significant three-way interaction finally indicated that genotypic differences during OF were dependent upon both, sex and trial (F (1,34) =15.06; p<0.0001) with male 5XFAD mice having the highest level of activity during OF1.

**Fig 2:**
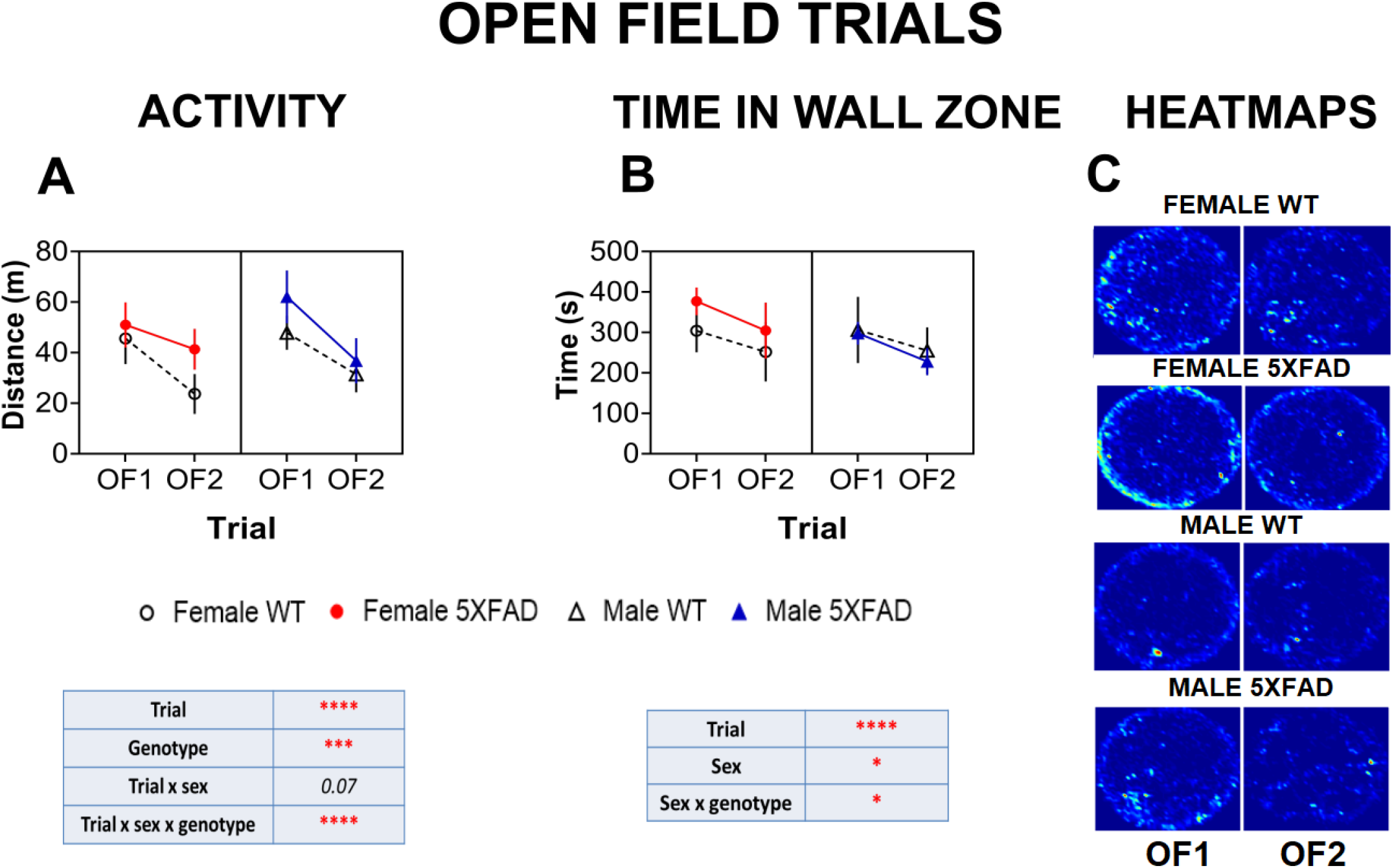
Activity measures for the open field trials (OF1 and OF2): Total distance travelled (in meters, m, per trial) during OF1 and OF2 trials for female and male 5XFAD vs WT mice (**A**), and total time spent in the thigmotaxis zone (in seconds, s) (**B**). Tables below graphs indicate statistically significant outcomes from a 3-way ANOVA with sex, genotype and trial (repeated measures) as factors. The pattern of exploratory behaviour for each group is also illustrated by representative heatmaps of one individual animal belonging to each group where bright colours indicate locations of frequent visits and darker colours indicate areas of rare approaches during OF1 and OF2 (**C**). Full statistical details can be found in Suppl. 1. All data are group means ±SD, Significances are indicated as: ****=p <0.0001, ***= p<0.001, *=p<0.05.

To observe if both *within-* and *between*-trial habituation occurred during the OF trials, 2-way ANOVAs for each sex and genotype group were conducted (binned per minute over OF1 or OF2, with trial and time as repeated measures). Results suggested both intact *within*- and *between-*trial habituation for all groups, yet the effect was (based on significances obtained) smallest for 5XFAD female mice (see Supplementary Figure S1 including statistics). Hence, 5XFAD mice differed in terms of activity, and to a lesser extent in the degree of habituation, in a sex-dependent manner.

#### Heightened thigmotaxic behaviour in the open field in female 5XFAD mice

Thigmotaxis, assessed as wall hugging (time in wall zone), was also affected by sex (F (1, 34) = 5.406; p<0.05, 3-way ANOVA, Fig. 2B). A strong effect of trial (OF2 vs. OF1), i.e. a decline between trial 1 and trial 2, was also noted (F (1, 34) = 46.92; p<0.0001). In contrast to activity, there was no overall genotype for thigmotaxis between 5XFAD and WT animals, yet inspection of data and an interaction obtained (sex x genotype: F (1, 34) = 6.190; p<0.05) indicated genotypic differences were present due to female 5XFAD animals displaying increased thigmotaxis compared to their WT counterparts. The representative heatmaps which indicated location preference for each trial for each genotype and sex also displayed higher occupancy of female 5XFAD mice in the wall zone (see heatmaps in Fig. 2C) especially during OF1.

#### Exploration of a novel object (NO) in 5XFAD mice

We next introduced an object (wire mesh cylinder) into the centre of the open field arena and returned test subjects for two further trials. Since global path length is not the best indicator for habituation during NO trials [42], object exploration and recognition was indexed by the time the animal spent in the interaction zone (Fig. 1C) and the number of visits to this zone.

No gross overall differences between 5XFAD animals and controls were observed for contact time in the interaction zone (Fig. 3A), but a main effect of sex emerged (F (1, 34) = 8.759; p<0.05), such that female animals spent less time in the zone compared to males. The lowest time in zone was recorded for female 5XFAD, in line with a trend for a sex by genotype interaction (F (1, 34) = 3.980; p=0.0541), suggesting subtle genotypic differences for female animals, especially during initial exploration of the object in NO1 (Fig. 3A). A strong overall effect of trial was also detected (F (1, 34) = 15.14; p<0.0001), although the horizontal nature of the graph for the female 5XFAD animals indicated a potential lack of between-trial habituation in these animals. Indeed, the familiarity index for time in zone suggested that time spent in trial 1 and trial 2 did not differ in female 5XFAD animals (Fig. 3B), but a mean value of the index close to 0 and even negative for some animals strongly suggests a deficit in object familiarity. In female WT animals, there was a robust trend (p=0.08) with regards to intact recognition. Both male 5XFAD and WT mice displayed significant recognition (p<0.05; one-sample t-test for each group against chance=0).

**Fig. 3:**
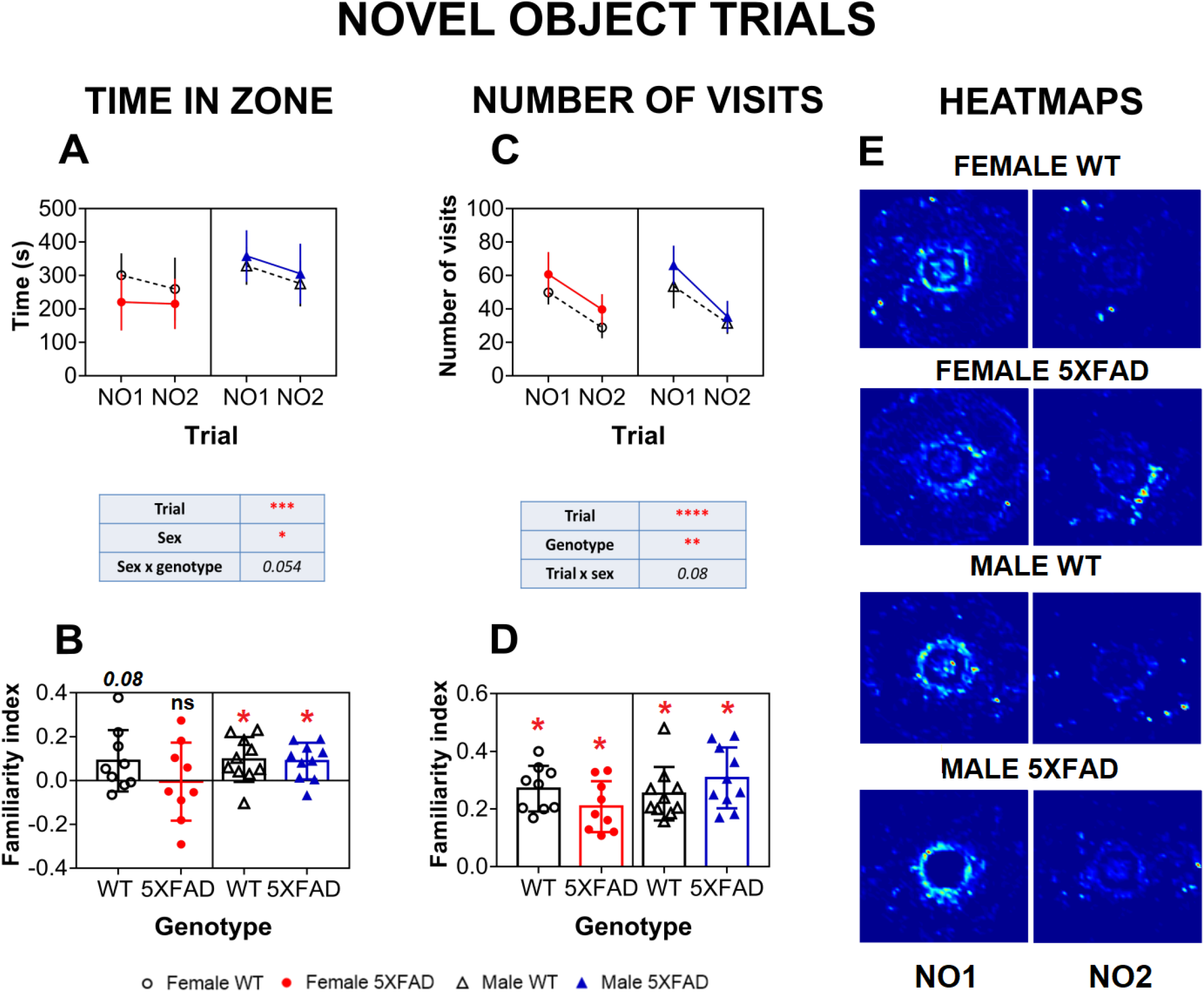
Exploration of the novel object during NO1 and NO2: Total time in the exploration zone (in seconds, s, per trial) during NO1 and NO2 for female and male 5XFAD vs WT mice (**A**) and total number of visits to this zone (**C**). Tables below graphs indicate statistically significant outcomes from a 3-way ANOVA with sex, genotype and trial (repeated measures) as factors. Familiarity indices in the form of scatter plots for time in zone **(B)** and number of visits **(D)** for female and male 5XFAD and WT mice. Results of one-sample t-tests for each group (against chance), and significances (or lack of) are indicated above the bars. No group difference using a one-way ANOVA was observed. Representative heatmaps for exploratory activity during NO trials (E) clearly reveal how pattern of exploratory activity has shifted to the novel object zone during these trials (brighter colour concentrated around the centre where object was placed). Full statistical details can be found in Suppl. 1. Data are presented as group means ± SD with symbols representing individuals; significances are indicated as: ****=p <0.0001, ***= p<0.001, *=p<0.05, ns=not significant.

As for the number of visits (Fig. 3C), 5XFAD mice entered the interaction zone more frequently than WT animals, as indicated by a strong effect of genotype (F (1,34) = 11.15; p<0.001). A robust trial-wise decrease in the number of visits was also observed in all groups (F (1,34) =232.2; p<0.0001), alongside a trend for a trial x sex interaction (F (1,34) = 3.064; p=0.08), indicative of trial effects affected by sex. The familiarity index for the number of visits to the zone (Fig. 3D) differed significantly (from the hypothetical value of 0) for all groups (p<0.05), with fewer visits made during the second trial.

5XFAD animals continued to display higher overall activity during the NO stage (genotype: F (1, 34) = 8.004; p=0.0078, see Supplementary Fig. S2), which was also dependent on the trial (trial: F (1, 34) = 317.9; p<0.0001). Heatmaps for representative animals of each group display both reduced time in the zone around the cylinder for female 5XFAD compared to other groups during NO1 and visibly reduced activity levels during NO2 vs NO1 for most groups except for the female 5XFAD animals (Fig. 3E).

#### Exploration during social interaction trials (SI) in 5XFAD mice

In the third and final test stage, we placed a sex-matched conspecific as a stranger into the cylinder and again recorded time in and entries into the interaction zone. During SI, a genotype difference for time in zone was obtained (F (1, 34) = 7.386; p<0.05); this was especially apparent for female 5XFAD animals during SI1 with the lowest contact time (Fig. 4A). A strong effect of trial was also observed (F (1, 34) = 18.69; p<0.0001) indicating between-trial habituation for all cohorts. Female 5XFAD animals displayed similar overall exploration times in both trials; this was confirmed by the lack of significant difference (against a value of 0) in their familiarity index (Fig. 4B), while all other groups (female WT, male 5XFAD and male WT) displayed positive, significant indices (p<0.05) for this parameter.

**Fig. 4:**
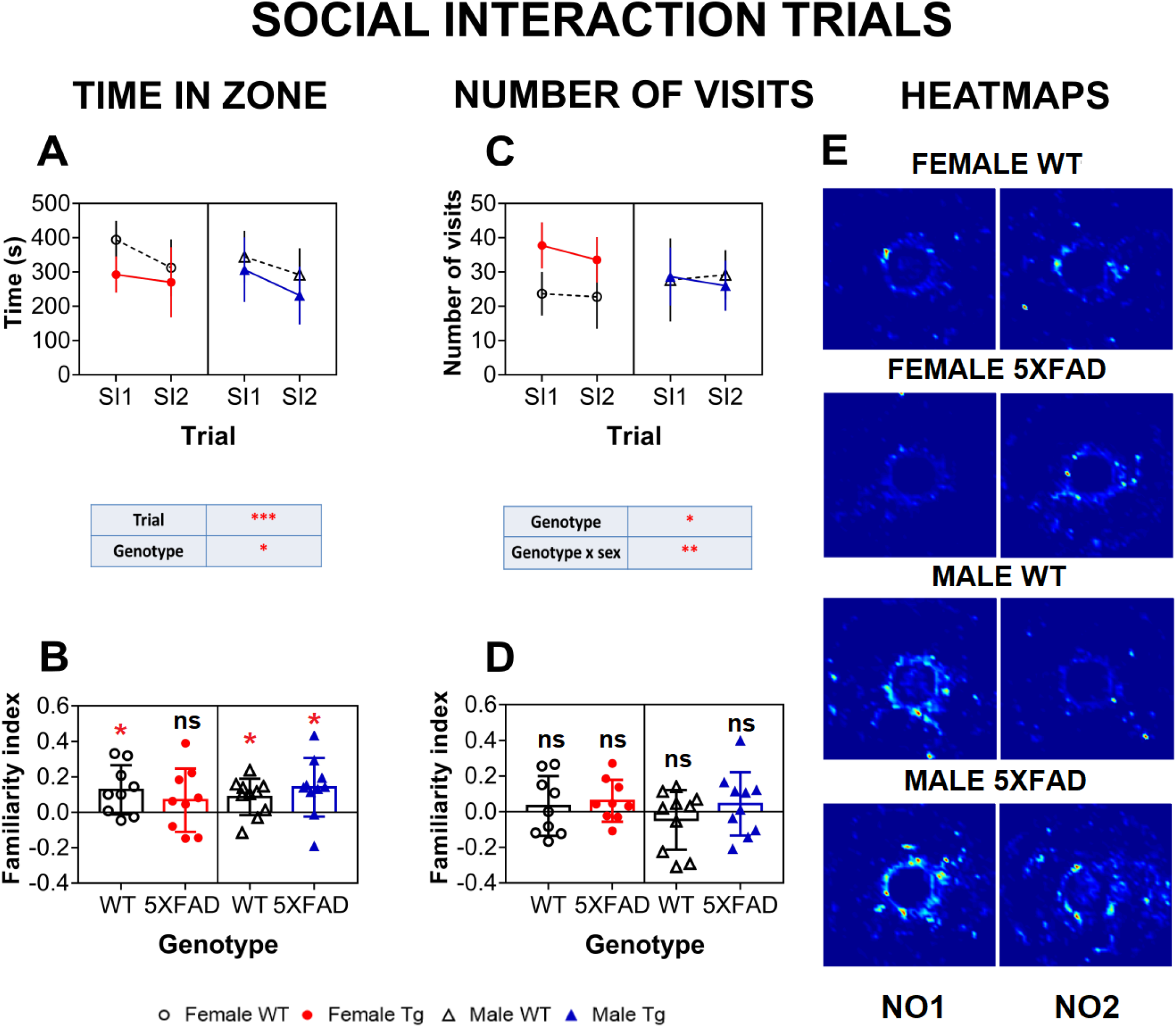
Exploration of the conspecific during SI1 and SI2: Total time in the exploration zone (in seconds, s, per trial) during SI1 and SI2 for female and male 5XFAD vs WT mice (A) and total number of visits to the zone (C). Tables below graphs indicate statistically significant outcomes from a 3-way ANOVA with sex, genotype and trial (repeated measures) as factors. Familiarity indices in the form of scatter plots for time in zone (B) and number of visits (D) for female and male 5XFAD and WT mice. Results of one-sample t-test done within each group (against 0), and significances (or lack of) are indicated above the bars. No group differences using a one-way ANOVA were observed. Heatmaps for exploratory activity (brighter colour marks more frequent visits) during SI trials (**E**) are displayed. Full statistical details can be found in Suppl. 1. Data are presented as group means ± SD, Significances are indicated as: ****=p <0.0001, ***= p<0.001, *=p<0.05, ns=not significant.

A difference between 5XFAD and WT animals (F (1, 34) = 6.299; p<0.05) was also obtained for the number of visits to the interaction zone (Fig. 4B) during SI trials; female 5XFAD mice visited the social partner more frequently than their WT counterparts during both SI trials (genotype x sex interaction: p<0.001). Together with contact time data, this suggests that female 5XFAD mice entered the zone more frequently, but visits were much shorter. Interestingly, there was no trial effect for number of visits (trial: p>0.05) during SI for any of the groups, likely due to the stronger social vs inanimate stimulus [42]. This was supported by the lack of significant familiarity index for the number of visits to the zone (p>0.05), a parameter that differentiates between object and stranger exploration.

During SI, the overall activity was not different between 5XFAD and WT animals (Supplementary Fig. S2). However, a trend for a sex difference emerged (F (1, 34) = 3.393; p=0.07), with female 5XFAD animals exhibiting sustained, heightened activity (sex x genotype interaction: (F (1,34) =3.383; p=0.08; sex x trial interaction: F (1, 34) = 6.665; p=0.0143). This is in line with the more frequent but shorter number of visits for female 5XFAD mice.

Collectively, the 5XFAD cohort, irrespective of sex, was robustly hyperactive during the OF-NO-SI paradigm and exhibited more frequent visits to the interaction zone during NO and SI trials. Gross object / stranger exploration differences between genotypes were detected for number of visits (NO and SI) and time in zone (SI only), while multiple interactions and the familiarity index supports a stronger deficit in females. Furthermore, during NO and SI, female 5XFAD subjects showed no *between-trial* habituation in exploration time but entered the interaction zone more frequently during the SI trials compared to all other groups. This is also further reflected in the heatmaps, which confirm that the female 5XFAD group spent less time in the vicinity of the stranger mouse, especially in SI1 with little change between trials (Fig. 4E).

### Post-mortem molecular analysis

#### Heightened gene expression of hAPP, but not hPSEN-1 in female 5XFAD mice

In line with the behavioural phenotype, 5XFAD female mice had significantly higher hAPP gene expression compared to males (p<0.05, +3% cf males; Fig. 5A). By contrast, gene expression of hPSEN1 was not different between sexes (Fig. 5B). As stated earlier, no hAPP and hPSEN1 was detected in WT animals.

**Fig. 5:**
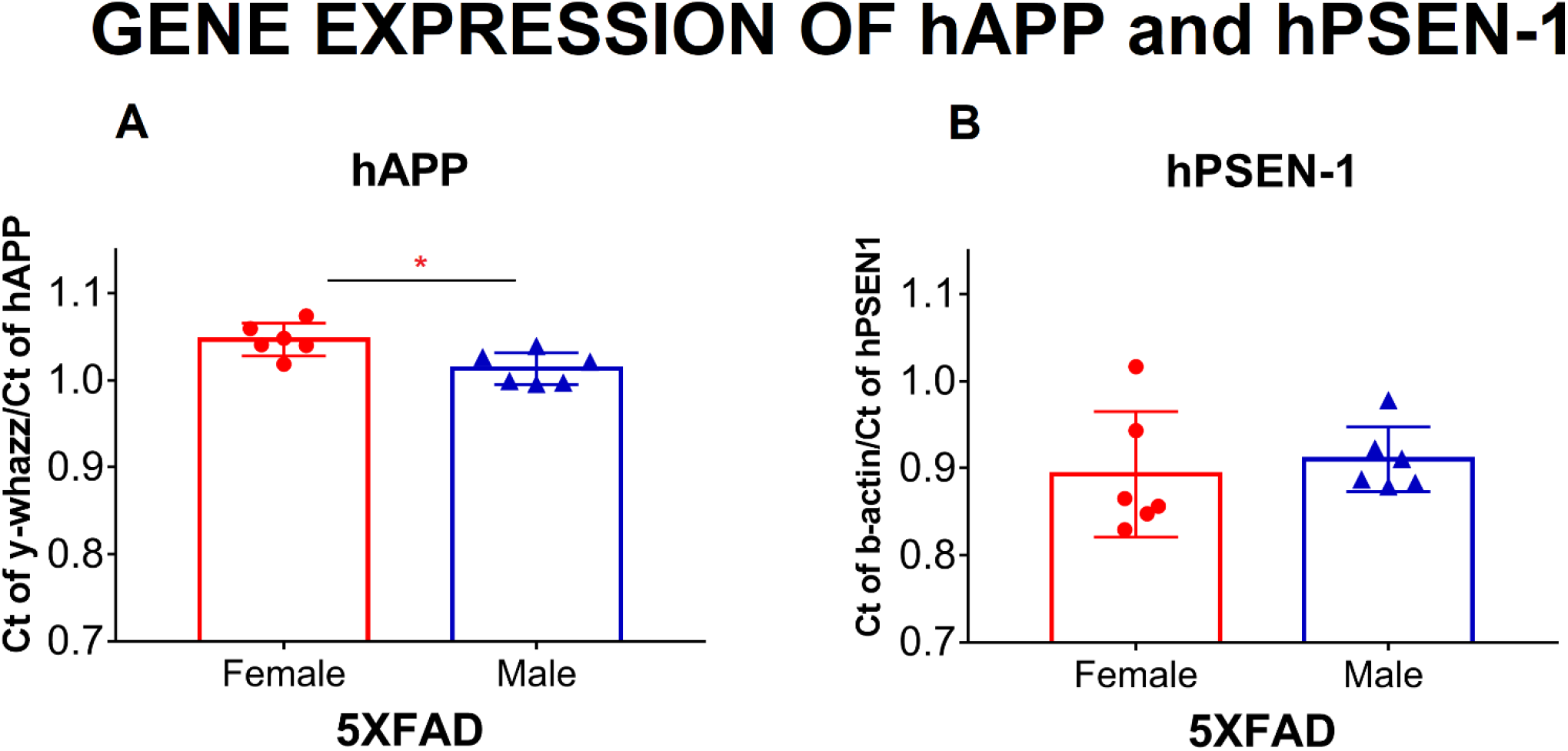
Gene expression of hAPP, but not PSEN-1 is higher in female 5XFAD animals compared to male transgenic mice: Gene expression of hAPP (**A**) and PSEN-1 (**B**) for 5XFAD animals of both sexes are reported as ratio of Ct values of most stable reference gene on plate (y-whazz/b-actin) with gene of interest (hAPP/hPSEN-1). Data are illustrated as scatter plots with mean ±SD; significance is displayed as: *=p<0.05 (parametric t-test).

#### Heightened levels of APP and its metabolites in female 5XFAD animals in soluble fraction

Full-length APP (fAPP) was determined by western blotting using the 6E10 antibody in both the *soluble fraction* (Fig. 6A) and *insoluble fraction* (Fig. 6B); results confirmed highly significant effects of genotype (*soluble*: F (1,27) =912.4, p<0.0001; *insoluble*: F (1,25) = 145.6, p <0.0001), sex (*soluble*: F (1,27) = 111.2, p<0.0001; *insoluble*: F (1,25) = 6.671, p=0.0160) as well as interactions (*soluble*: F (1,27) = 111.2, p<0.0001; *insoluble*: F(1, 25) = 4.698, p=0.039). Compared to very low levels in WT mice, significantly elevated amounts of fAPP were found in female 5XFAD animals, also compared to male 5XFAD and particularly in the soluble fraction (+100% for *soluble,* +45% for *insoluble; soluble*: p<0.0001; *insoluble*: p=0.01) (Fig. 6A and 6B).

**Fig. 6:**
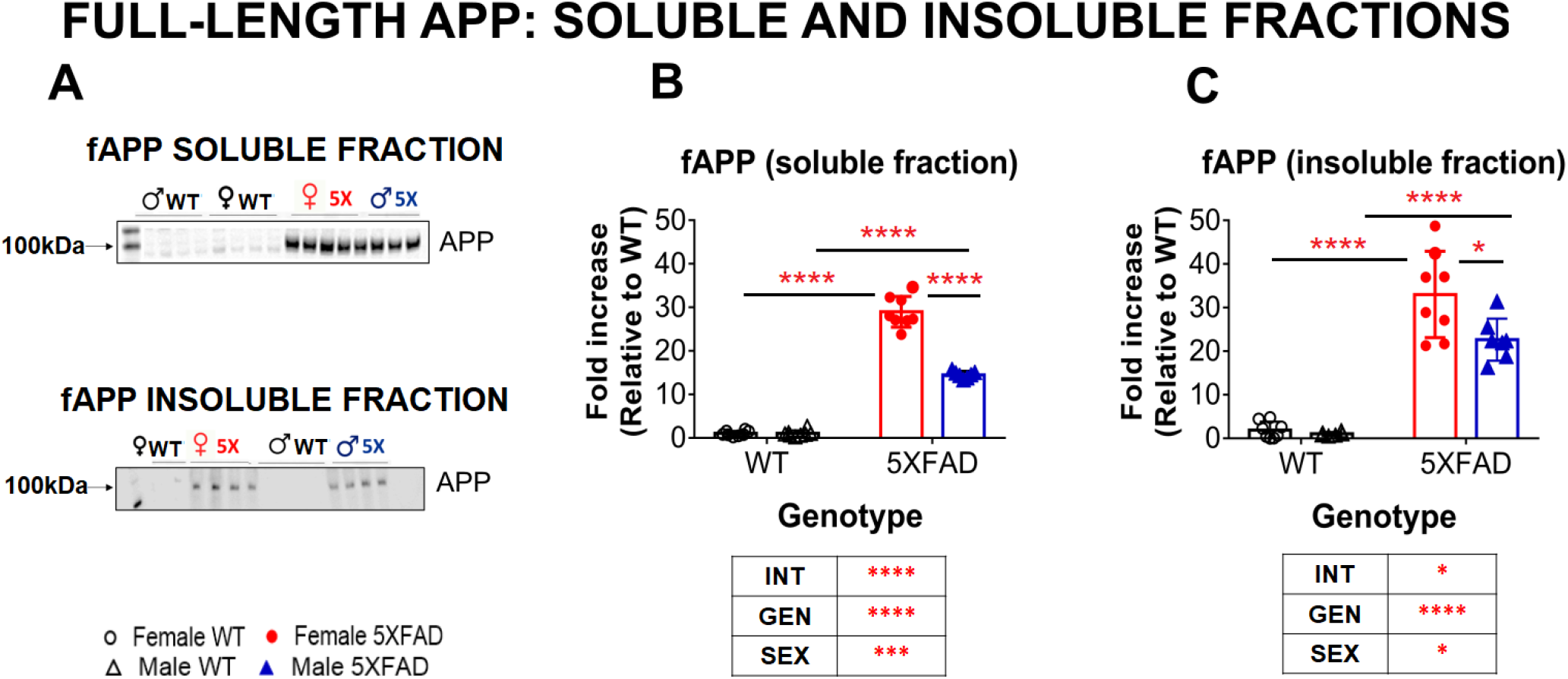
Greater levels of full-length APP in female 5XFAD mice in soluble and insoluble fractions: Representative images of western blots of both soluble and insoluble fractions probed with antibody 6E10 for the detection of full-length APP (fAPP) with molecular weight in kDA (**A**). Protein levels were higher in female 5XFAD compared to male transgenic animals in both the soluble (**B**) and insoluble (**C**) fractions. All data were normalised to female WT with data sets visualised as scatter plots with mean ± SD; tables indicate significant 2-way ANOVA effects where int=interaction, gen=genotype; significances for individual genotypes/sexes are displayed on graph as ****=p<0.0001, **=p<0.01 (Bonferroni post-hoc test).

Dot-blot analysis of Aβ protein was carried out using MOAB-2 antibody, which does not cross-react with APP or non-Aβ metabolites and has lower affinity for Aβ40 compared to the more neurotoxic Aβ42 (see [45]). In the *soluble* fraction (Fig. 7B), data mirrored results obtained with 6E10, in that we found a very strong effect of genotype (F (1, 28) = 98.41, p<0.0001), sex (F (1,28) = 17.07, p=0.0003) and a genotype x sex interaction (F (1, 28) = 12.73, p=0.0013), with higher levels of Aβ in female 5XFAD mice compared to their male counterparts (+108% cf male 5XFAD; p<0.0001). An effect of genotype was also observed in the *insoluble* fraction (Fig. 7C; (F (1, 25) = 12.85, p=0.0014)) even though values were generally lower and more variable, which may explain why the effect of sex did not reach statistical significance (Fig. 7C).

**Fig. 7:**
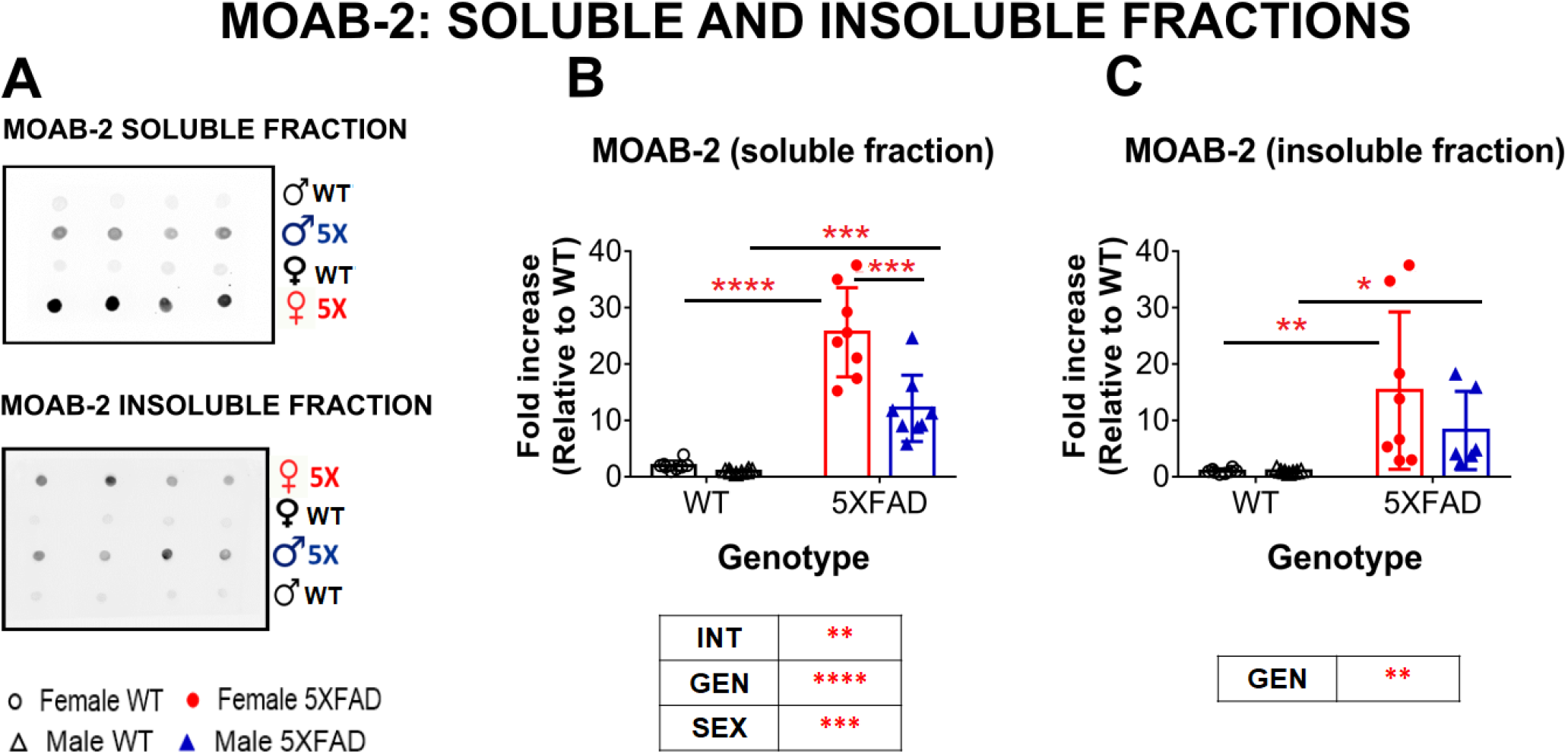
Heightened MOAB-2 levels in female 5XFAD mice compared to male 5XFAD mice in soluble but not insoluble fractions: Representative images of dot blots of both soluble and insoluble fractions probed with MOAB-2 for the detection of Aβ (**A**). Protein levels were higher in female 5XFAD animals compared to male transgenic animals in soluble (**B**) but not insoluble (**C**) fraction. All data were normalised to female WT and visualised as scatter plots with mean ± SD; tables indicate significant 2- way ANOVA effects where int=interaction, gen=genotype; significances for individual genotypes/sexes are displayed on graph as ****=p<0.0001, ***=p<0.001, **=p<0.01 (Bonferroni post-hoc test).

Overall, both gene and protein analyses confirmed hAPP expression and processing in 5XFAD mice, with a higher gene and protein load for female vs male 5XFAD mice confirming previous reports [31]. Protein levels were even more dramatically increased in females than suggested by gene expression, indicative of additional factors influencing pathology build-up.

#### Increased inflammation in female 5XFAD mice

Western blot analysis of the astrocyte marker GFAP indicated a significant overall effect of genotype (F (1,25) = 13.07, p=0.001), yet significantly higher levels of GFAP were only found in female 5XFAD animals compared to female WT (+150%, p=0.047; Fig. 8B). Immunoblot analysis of the activated microglial marker, Iba-1 (Fig. 8C), indicated a strong significant effect of genotype (F (1,26) = 17.53, p=0.003) but also sex (F (1,26) =16.95, p=0.003). Post-hoc analysis of Iba-1 confirmed that female 5XFAD mice had higher levels compared to female WT (+50%, p=0.002), but also vs. male 5XFAD mice (+50%, p=0.0018). A lack of specificity observed while trying various NLRP3 antibodies was replaced by utilising a qPCR-based gene expression approach to analyse this inflammasomal marker (Fig. 8D). It also indicated a significant effect of genotype (F (1,17) = 22.08, p=0.0002) and sex (F (1,17) = 8.956, p=0.008) with female 5XFAD animals yielding higher expression levels of NLRP3 compared to male 5XFAD (+64%, p=0.03) and to their respective WT (+130%, p=0.001). Here, no genotype effect was observed for the male mice.

**Fig. 8:**
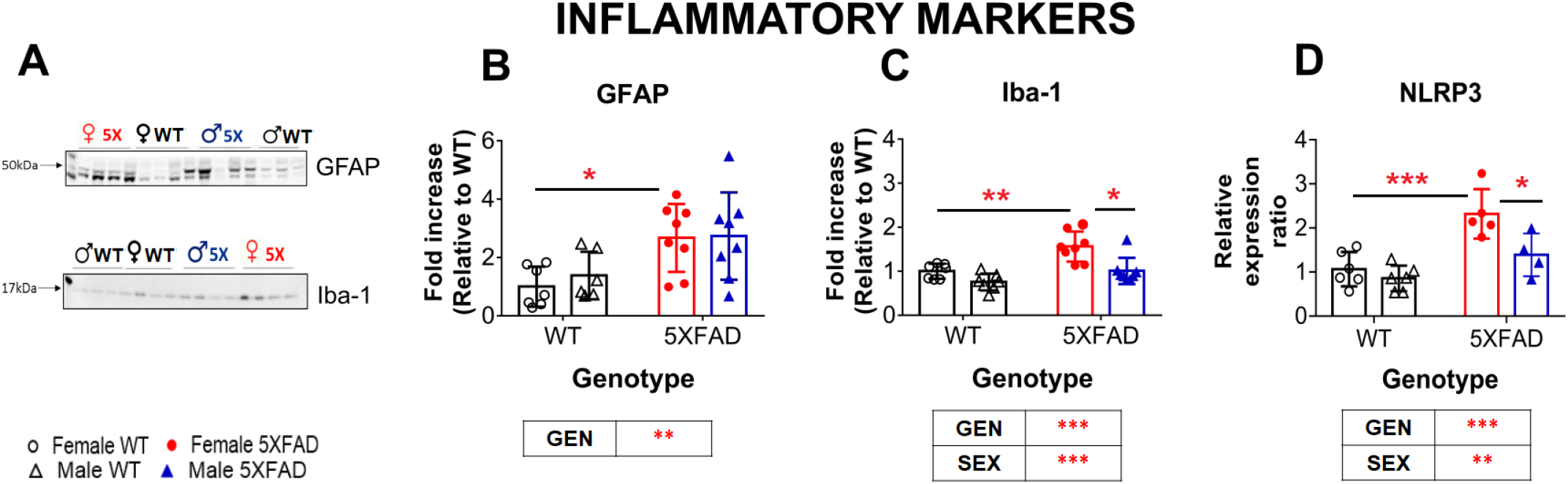
Inflammation was enhanced in 5XFAD mice: Representative images of western blots for GFAP and Iba-1 with corresponding molecular weights in kDa (**A**). Group average protein levels for GFAP (**B**), Iba-1 (**C**), and gene expression of NLRP3 (**D**) were higher in female 5XFAD animals compared to their corresponding WT and compared to the male 5XFAD (for Iba-1 and NLRP3). All data were normalised to female WT and visualised as scatter plots with mean ± SD; tables indicate significant 2-way ANOVA effects where int=interaction, gen=genotype; significances for individual genotypes/sexes are displayed on graph as, ***=p<0.001, **=p<0.01, *=p<0.05 (Bonferroni post-hoc test).

#### No changes in synaptic markers in 5XFAD mice

Protein levels of the presynaptic marker synaptophysin revealed no overall difference between genotypes; however, a sex effect was detected (F (1, 25) = 6.488, p=0.017; Fig. 9B), due to higher levels observed in females in both genotypes. Immunoblot analysis of the post-synaptic marker PSD-95 (Postsynaptic density protein 95; Fig. 9C), on the other hand, indicated an effect of genotype (F (1,26) = 5.174, p=0.03), with increased levels in transgenic mice. It therefore appears that despite the high levels of amyloid pathology, a gross reduction of synaptic markers could not be confirmed in both sexes of 5XFAD mice.

**Fig. 9:**
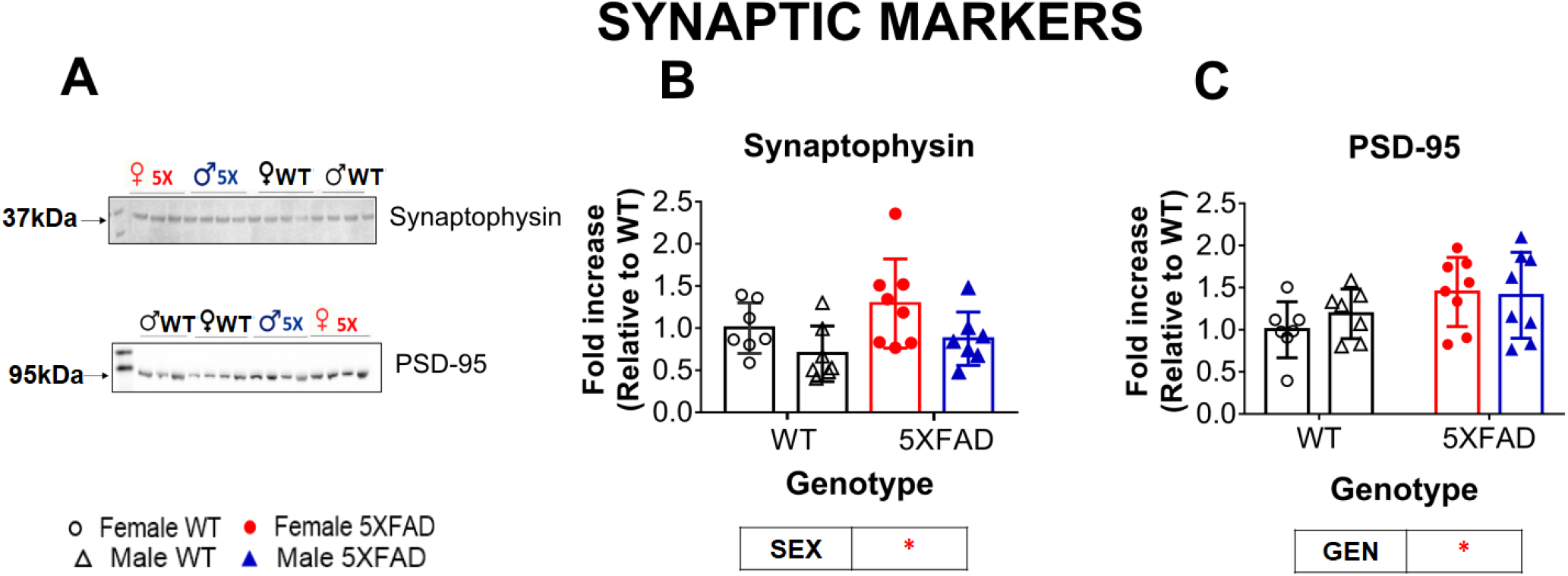
No major changes in pre- and post-synaptic markers: Representative images of Western blots for synaptophysin and PSD-95 with corresponding molecular weights in kDa (**A**). A subtle sex effect (but no effect of genotype) was noted for synaptophysin such that females had heightened levels of this presynaptic protein (**B**). Protein levels of PSD-95 were higher in 5XFAD animals only relative to their WT (**C**). All data were normalised to female WT and are presented as scatter plots with mean ± SD; tables indicate significant 2-way ANOVA effects where int=interaction, gen=genotype; significances for individual genotypes/sexes are displayed on graph as *=p<0.05 (Bonferroni post-hoc test).

#### Sex-associated differences in specific arms of the ER pathway

Gene expression analyses of different markers from the ER stress pathway was conducted next (Fig. 10). In the event of chronic ER stress in AD, the chaperone, BiP (Binding immunoglobulin Protein), which supports folding of proteins, can cause the activation of the UPR (Unfolded Protein Response) signalling and transcription factors, IRE1 (inositol requiring enzyme), ATF6 (activated transcription factor 6), CHOP (C/EBP Homologous protein) and total XBP (XBPt or X-box protein, a further key transcription factor involved in ER degradation) which forms spliced XBPs (XBPs).

**Fig. 10:**
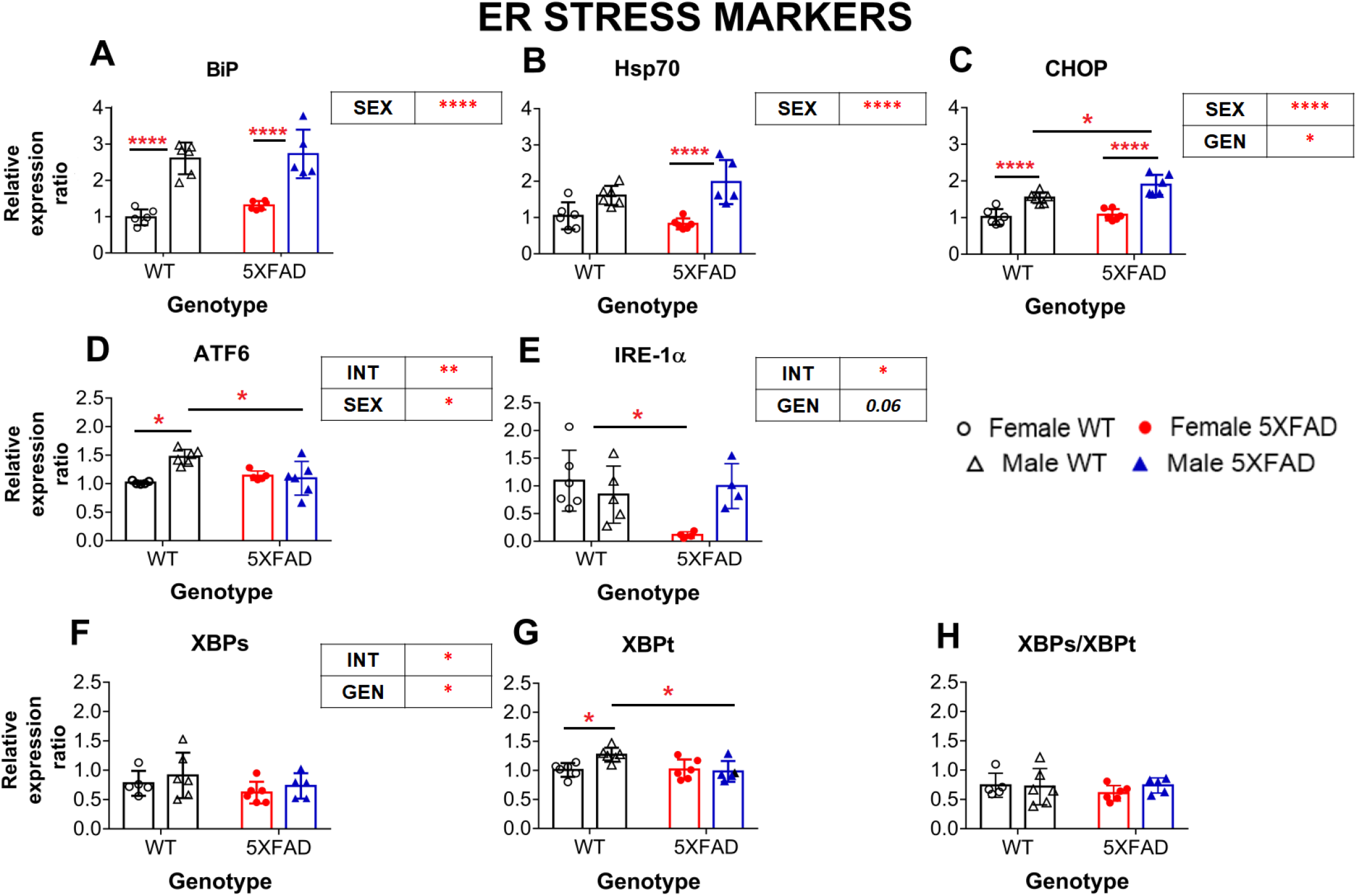
Heightened expression of HSP-related markers in male mice: Gene expression of ER stress markers BiP (**A**), Hsp70 (**B**), and CHOP (**C**) were higher in male 5XFAD animals compared to female 5XFAD. Male WT animals also had higher expression compared to female WT animals in case of BiP and CHOP. In addition, male WT had higher expression of ATF6 (**D**) and XBPt **(F)** compared to both male 5XFAD and female WT animals. Female 5XFAD animals had very low expression of IRE1α expression (**E**) compared to female WT and male 5XFAD (trend) while neither XBPs (**G**) nor the ratio of XBPs/XBPt (**H**) revealed any differences. All data are presented as scatter plots with mean ± SD; tables indicate significant 2-way ANOVA effects where int=interaction, gen=genotype; significances for individual genotypes/sexes are displayed on graph as, ****=p<0.0001, *=p<0.05 (Bonferroni post-hoc test).

Following on from the above tissue results, gross genotype effects were only observed for 2 markers: CHOP (F (1,20) =5.96, p=0.024; Fig. 10B) and total XBP (XBPt) (F (1,19) = 5, p=0.037; Fig. 10F). However, while male 5XFAD animals had higher expression of CHOP (p=0.046, Bonferroni’s post-hoc; +50%), they had lower expression of XBPt (p=0.028; −25%) compared to male WT.

In addition, sex-dependent differences were detected for the ER chaperone BiP (F (1, 19)= 82.14, p<0.0001; Fig. 10A), as well as the global chaperone marker Hsp70 (Fig. 10B; F(1, 19)=30.31, p<0.0001). CHOP, a transcription factor involved in ER-associated apoptosis regulation (F(1, 20)=64.34, p<0.0001; Fig. 10C) followed the same pattern.

In stark contrast to amyloid and inflammation markers, male 5XFAD animals had higher expression of BiP (p<0.0001; +100%), CHOP (p<0.0001; +100%) and Hsp70 (p=0.002; +120%; Fig. 10A-C) compared to female transgenic animals. For both CHOP and BiP, male WTs also showed higher expression compared to the female WTs (p<0.0001, p=0.0015, respectively).

An interaction of sex x genotype was noted for gene expression of ATF6 (F(1,18) =11, p=0.0038; Fig 10D) which also reported an effect of sex (F(1, 18)=7.524, p=0.0134). A matching pattern (interaction of sex and genotype) was also observed in the case of XBPt (F (1,19) =5.614, p=0.028, Fig. 10F). For both ATF6 and XBPt, male WT animals had higher gene expression compared to both male 5XFAD (ATF6: p=0.009; +50%, for XBPt see above) and female WT (ATF6: p=0.003, +50%; XBPt: p=0.039, +25%) animals. Neither the spliced variant of XBP (XBPs) nor the ratio of XBPs/XBPt displayed any effect of genotype or sex (Fig. 10G and 10H)).

An unusual sex x genotype interaction was also uncovered for IRE1α (F (1,15)=7.254, p=0.016, Fig. 10E), alongside a trend for a genotype effect (p=0.07), with female 5XFAD animals reporting very low expression (cf WT: p=0.026, −91%;, cf male 5XFAD: p=0.08, −78%). No such genotypic difference was observed for males. Overall, ER stress markers by and large yielded sex-specific and UPR-arm specific alterations, also causing a number of sex-dependent genotype effects.

#### Heightened levels of neurotrophic markers in male mice

The downregulation of neurotrophic factors such as BDNF and its TrkB receptor, as well as CREB, which controls the transcription of BDNF, have all been reported in AD, and considered as targets for treatment [48]. At the same time, sex differences in the expression of neurotrophic factors have also been well documented [49], making these protective entities interesting and relevant markers for the present study.

Analysis of the gene expression of BDNF (Fig. 11A) did not yield any reliable differences between the sexes or genotypes (but note trend for genotype: F (1, 18) =3.611, p=0.076). For CREB and TrkB, a significant effect of sex was seen (F (1,16) = 46.46, p<0.0001 and F (1,19) = 10.89, p=0.004, respectively). Post-hoc analysis of CREB (Fig. 11B) suggested higher expression in males cf. females for both genotypes (p=0.0028, +66% for 5XFAD; p=0.0005, +95% for WT), while for TrkB (Fig. 11C), male WTs yielded higher levels compared to male 5XFAD (p=0.048, +36%) and female WT animals (p=0.0006, +50%). It therefore seems that, like ER stress markers, male mice express enhanced levels of neurotrophic and potentially protective factors, particularly CREB.

**Fig. 11:**
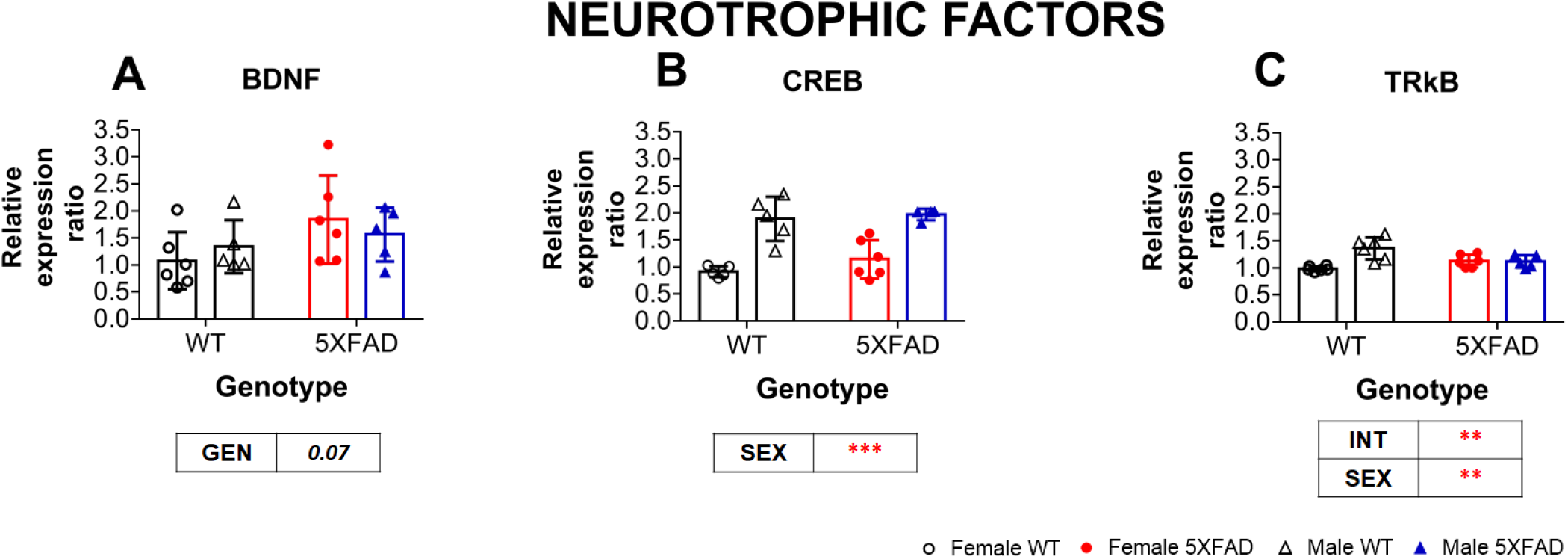
Heightened neurotrophic factor expression in male mice: Gene expression of neurotrophic factors revealed no genotypic or sex differences in BDNF (**A**). CREB (**B**) was higher in male 5XFAD (or WT) compared to female 5XFAD (or WT) but TrkB was found to be higher in male WT compared to female WT as well as male 5XFAD animals (**C**). Data are presented as scatter plots with mean ± SD; tables indicate significant 2-way ANOVA effects where int=interaction, gen=genotype; significances for individual genotypes/sexes are displayed on graph as ***=p<0.001, **=p<0.01, *=p<0.05 (Bonferroni post-hoc test).

### Correlational analyses

To make advancements regarding our understanding of links between pathways and phenotypes, and to explore connections between cognition, AD pathology and related cellular markers, we next conducted a multi-factorial correlation analysis in both WT and 5XFAD animals (Fig. 12). The greatest power for this analysis is achieved by combination of data sets derived from male and female cohorts. Additional sex-specific heatmaps provided (Fig. 12 B, C,E,F) are for comparison only; we did not analyse those further to avoid type 1 and type 2 errors. Particular weight is placed on the emergence of clusters comparing i) behavioural proxies; ii) cellular proxies including pathological and inflammatory, but also stress and trophic markers; and iii) behavioural x cellular proxies and their differences *between* genotypes.

**Fig. 12:**
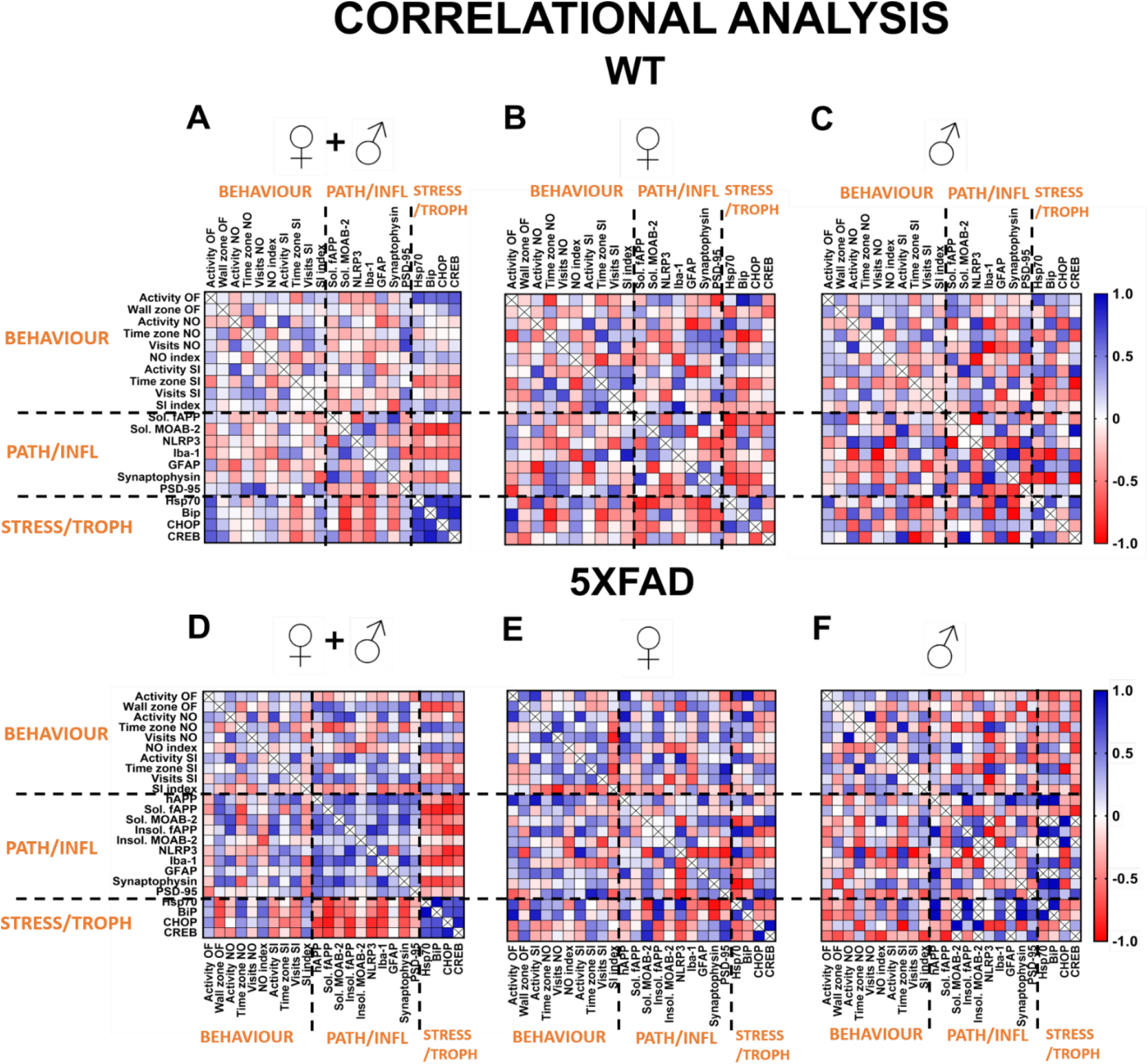
Correlations matrices between behavioural and molecular markers: Heat plots depict Z-score converted (overall mean) correlation matrices with positive (blue) and negative (red) correlations based on Pearson’s r correlational coefficient, for WT ♀+♂ (**A**), WT♀ (**B**), WT ♂ (**C**) 5XFAD ♀+♂ (**D**), 5XFAD ♀ (**E**), 5XFAD ♂ (**F**) animals. Behavioural and tissue markers are placed in sequence of their examination. Upper and lower triangle represent mirror images and dashed lines denote proxy categories (behavioural, pathology and inflammation, stress and trophic support. Abbreviations utilised are as follows: Activity OF/NO/SI= total activity in OF1+OF2/NO1+NO2/SI1+SI2; wall zone OF= total time in wall zone during OF; Time zone NO/SI= total time in interaction zone during NO (NO1+NO2)/SI (SI1+SI2); Visits NO/SI= Total number of visits during NO (NO1+NO2)/SI(SI1+SI2); NO/SI index= familiarity index for time in zone during NO or SI; Sol. = Soluble ; insol.= insoluble; Path/infl= pathology/inflammation indicating clusters formed by markers for amyloid, synaptic or inflammatory pathways; Stress/Trophic= ER stress, neurotrophic indicating clusters formed by markers belonging to ER stress/neurotrophic pathways. For further detail, see Results.

#### Correlations between behavioural proxies

Overall, only modest correlations between the different behavioural proxies were apparent, and this was similar in WT and 5XFAD cohorts. The strongest positive correlations (blue fields in Fig. 12A) were between activity measures of all stages of the task and time spent in the interaction zone for SI and NO stages, suggesting that animals with heightened locomotor activity also more readily visited the object/social partners. This was observed in both genotypes and is in line with a lack of an overt cognitive impairment in 5XFAD mice. In extension to our previous work using this paradigm [42], there was no correlation between the overall activity during OF and any other stage of testing in WT mice; by contrast, ambulatory activity correlated positively between all stages of testing in 5XFAD mice (blue squares in first row, Fig.12A, D) strengthening our argument of the predominant activity-related phenotype in this AD model. Intriguing was also the lack of clear correlations between the familiarity indices and time in contact zone or number of visits to these zones. While reasons for this are unknown, it was observed in both study cohorts suggesting that the familiarity index is differentially influenced by multiple behavioural parameters.

#### Correlations between cellular markers

Heatmaps differ between genotypes for molecules associated with cellular pathology. This is mainly due to the specific expression of human APP and the resulting amyloid species detected in 5XFAD mice, and which are absent in WT. Nevertheless, some clear clusters arose from the comparisons, more clearly seen in 5XFAD tissue. Correlations were globally positive between amyloid, inflammatory and synaptic markers in 5XFAD brains (Fig. 12D), some of them highly significant (dark blue squares) clearly underlining their relationships, especially on the background of tissue stress caused by amyloid levels. This was not as clear-cut in WT mice, a more varied pattern between mildly positive (faint blue) and mildly negative (faint red) correlations emerged. Similar for both genotypes, however, was the observation that stress and neurotrophic markers correlated strongly with each other, but negatively with amyloid load, inflammation, and synaptic protein levels; some of these negative correlations were highly reliable (dark red squares).

#### Correlations between behavioural proxies and pathological molecules

Less homogenous were correlations between behavioural readouts and molecular endpoints. An overall scattered and lightly coloured field showed few consistent correlations. Of interest, especially in light of the heightened activity found in 5XFAD mice, are positive correlations (blue squares) between activity-related endpoints (time in wall zone, activity in NO and SI as well as visits to the object and social partner) and markers of amyloid, inflammation (Iba-1) and particularly presynaptic labelling, which were not found to the same degree in WT mice. Intriguingly, these correlations were even stronger in female 5XFAD mice (Fig. 12E) relative to males and may underpin the more pronounced phenotype in female transgenic animals. Finally, a strong correlation appeared between stress/neurotrophic markers and the global activity in the OF stage in WT mice (see dark blue raster, bottom of row 1, Fig. 12A). This characteristic was less pronounced in 5XFAD tissue (Fig. 12D), mainly due to male mice (Fig. 12F), which showed negative correlations for these same rasters. This would be in line with the proposed sex-specific protective or compensatory molecular response. Finally, both phenotypes showed positive correlations between familiarity indices for NO/SI vs. cellular stress/trophic markers. Given these indices offer a cognitive proxy of the subjects, this underlines the observation that no overt cognitive impairment was observed in 5XFAD mice.

## DISCUSSION

Our results indicate distinct and dissociable sex-biases in development of both behavioural and molecular pathology in a cohort of 5-month-old 5XFAD mice. Mirroring what has been frequently observed in human studies, sex differences in transgenic mouse models of AD suggest a greater female-specific propensity for enhanced pathology and AD related cognitive phenotypes [8,33,50,51]. Overall, our data indicate that female transgenic 5XFAD animals present with hyperactivity, heightened anxiety, and some subtle anomalies in object processing and social interaction in the OF-NO-SI paradigm, which were not found in males. They also exhibited higher levels of inflammatory markers and amyloid pathology, alongside reduced levels of protective neurotrophic or heat shock protein (HSP)-related markers. Furthermore, our correlation analyses in both WT and transgenic animals strongly suggests that the critical factors are not restricted to beta-amyloid itself, but rather dependent on the extent of the inflammatory reaction it causes, and how effective the compensatory strategies in the form of ER stress and neurotrophic factors are.

The behavioural profile obtained via OF-NO-SI testing [42] offered multiple within-subject readouts of behaviour, including activity and habituation, as well as interaction and recognition of an inanimate object vs. a conspecific. 5XFAD animals exhibited higher locomotor activity compared to WT during the OF trials. Both intact within- and between-trial habituation of activity was observed across all groups; yet the degree of habituation was least pronounced in female 5XFAD mice compared to all other groups, suggesting that previously reported deficits in habituation to an open-field apparatus observed in (mixed-sex) 5XFAD cohorts [38, 52] may also have been sex-specific. In partial agreement with our data, Leary et al. also reported that female 5XFAD animals (on a B6SJL background, unlike the C57BL/6 background used here) travelled more than their respective WT (at 6 months of age), yet no such difference was observed in the male cohort [35]. The relationship between heightened ambulation and thigmotaxic (anxiety) behaviour was also increased in female 5XFAD animals compared to WT [36]. Hyperactivity is common in AD models and has also been detected in other APP over-expression lines [53], while for example low-expression knock-in transgenic mice showed lower activity at 5 months of age [12]. In general, a positive link between activity and cognitive phenotypes has been previously suggested [54], but hyperactivity can also lead to a more disrupted exploration pattern (as seen here in the female 5XFAD mice), and disturbed circadian activity, as often reported in transgenic AD lines (reviewed in [55]).

We also observed that abnormalities in object exploration and social interaction behaviour are modulated by both genotype and sex. During NO, 5XFAD animals explored the novel object less compared to WT, especially during initial exploration of the object in NO1. Both, a lack of deficits in male [56] and in female 5XFAD mice have been noted previously for novel object exploration in 8-month old cohorts [57], but subtle sex-dependent phenotypes occurred in older subjects at 12 months [33]. In similar studies using TASTPM mice with mutant APPswe and PSEN1, both sexes displayed equal impairments on object exploration despite much higher Aβ load in females [58]. Although we here report a strong correlation of performance with amyloid load specifically affecting female 5XFAD mice, it appears that this relationship may be more complex and depend on the amyloid and non-amyloid factors [59].

Genotypic differences during SI were more pronounced (vs NO), with female 5XFAD animals spending less time in zone compared to their controls and displaying lack of between-trial habituation as assessed by the SI familiarity index for time in zone. To note, 12 month old transgenic 5XFAD female mice have previously exhibited less investigation of a novel stranger in a three-chamber apparatus as well as in a free social interaction test [39]. Given the greater anxiety-like phenotype in female 5XFAD animals, this could be indicative of novelty aversion due to heightened social anxiety. In other APP and PSEN1 mice, males at 6 months of age also displayed reduced sociability, again combined with hyperactivity [60]. Overall, transgenic mice with high levels of mutated APP and PSEN1 seem to consistently display changes in activity alongside reduced social interaction behaviours, with an additional dependency on sex.

We also found levels of both fAPP (soluble and insoluble fractions) and MOAB-2 reactive amyloid species (soluble fraction) to be higher in female 5XFAD animals compared to males as reported previously [31,32,61]. None of these studies, however, discriminated between full-length APP (and Aβ) pathology in different fractions. Given the importance of soluble pre-fibrillar Aβ species in tracking disease progression and decline in AD [45, 46], our results further confirmed higher soluble amyloid load in females compared to males. This had previously been attributed to higher hAPP expression [31, 32]; yet here, the increase in expression was very modest (+3% vs males), but amyloid-related protein levels were about double. Thus, gene expression alone is unlikely to fully account for the heightened amyloid accumulation. Moreover, no sex difference was observed for PSEN-1 gene expression, in line with a previous observation [32], though gamma-secretase activity may of course be higher in females. In any case, only one of the introduced transgenes is affected by sex, while the other is not. Congruent with the amyloid pathology, inflammatory markers GFAP and Iba-1 as well as NLRP3 were generally highest in the female 5XFAD group. The NLRP3 inflammasome has been implicated in AD and suggested to contribute to the pathology in APP/PSEN1 mice [62] by potentially acting as a sensor of Aβ, and by impairing microglial clearance and promoting inflammation [63, 64]. One previous report in 5XFAD mice found higher levels of NLRP3 in transgenic animals compared to WT; however this did not reach statistical significance [65], potentially due to the use of only male mice.

Our *within-subject* correlational analysis further confirmed the direct link between amyloid markers and inflammatory pathways in both control and 5XFAD mice, with stronger positive clusters observed in the transgenic animals. Positive correlations between activity measures (and number of visits) during NO/SI and both soluble and insoluble amyloid markers as well as inflammatory markers (in form of Iba-1) predominantly in the transgenic group strongly suggest that high levels of amyloid are causally linked with the hyperactivity/anxiety phenotype. As a corollary, familiarity indices correlated negatively with these pathology markers in 5XFAD mice.

Somewhat surprising was the unexpectedly higher (rather than lower) levels of the synaptic marker PSD-95 in 5XFAD mice. It is important to note that total protein levels were measured in our study rather than functional synapses (which would be more accurate). Changes in the level of synaptic markers depend largely on the region investigated and may be a late-occurring event in the pathology of AD [66]. The increase in PSD-95 levels has also been observed in 5XFAD females by others [33], who attributed it to the loss of PSD-95 from apical dendrites and their movement and build-up in cell bodies [67], but this remains unexplained functionally. We here show that more positive correlations (blue) between PSD-95 and behavioural proxies in WT mice turn into no or negative correlations (red) in 5XFAD subjects indicating that enhanced levels of PSD-95 is functionally detrimental. By contrast, the presynaptic marker synaptophysin was unchanged in 5XFAD mice but correlations with activity-dependent behavioural endpoints were generally positive, and with familiarity indices negative. This is more like the pattern found for pathology/inflammation markers and confirms that both pre-and postsynaptic alterations may be responsible for functional changes recorded in the OF-NO-SI. While Maiti and colleagues have confirmed earlier reports of reduced synaptophysin and PSD-95 in cortex and hippocampus in 5XFAD mice from 4-12 months of age [28,68–70], others have published data reporting no differences in both markers [71]. However, most studies used exclusively males from a mixed B6/SJL background and control mice often were purchased from different vendors.

ER stress and neurotrophic pathways in both the 5XFAD and WT groups revealed negative correlations with soluble and insoluble amyloid markers, indicating that reduced levels of chaperons (BiP, Hsp70 and CHOP) and neurotrophic factors associate with amyloidosis. In fact, the elevated levels of some ER stress markers and neurotrophic factors observed in males may have provided protection against amyloid-induced toxicity. ER markers all exhibited strong sex-dependent effects with male animals displaying much higher levels compared to females. It has been demonstrated previously that expression of HSPA1 and HSPB1 (which constitute Hsp70) along with six other heat-shock proteins (HSPs) showed male-biased expression in the hippocampus of both 5XFAD and their C57BL/6 WT controls at 4 months of age [32, 72]. HSPs in general stabilise proteins and prevent misfolding [73], while secreted Hsp70 masks Aβ in the extracellular space and suppresses neurotoxicity [74–76]. From this perspective, the higher expression of Hsp70 (and by extension BiP, a member of the Hsp70 family localised to the ER) in male mice may have offered some protection against Aβ accumulation and disease progression. Protein levels of CHOP were also increased in male APP/PSEN1 mice compared to their WTs [77], but our results are at odds with the lack of differences between genotypes and/or sex in protein levels of CHOP or BiP reported in 4-month 5XFAD mice previously [29]. Markers belonging to other arms of the UPR pathway such as ATF6 also displayed strong sex-related and genotypic differences (male WT>male 5XFAD). Investigations into the ATF6 pathway in transgenic models of AD remain limited [23] and to our knowledge this is the first report into its role in both sexes of the 5XFAD model. Of interest are also low levels of gene expression of IRE-1α in female 5XFAD animals and the lack of changes in expression of activated XBP (XBPs), which may reflect an age-dependent change in the directionality of gene expression of these markers [78]. Strong sex-dependent effects were also observed for neurotrophic markers like CREB and TrkB such that all male animals exhibited higher levels, yet here no genotype differences emerged. Similar to our observations, sex-related differences in CREB signalling (lower in females) have been described in 12 month old 3xTg-AD mice [79]. No strong positive or negative correlations between behavioural and ER stress/trophic markers were revealed. All correlations were moderate to weak and the scatter in WT is similar to the scatter in 5XFAD mice. Exception to this rule was the familiarity index for both NO and SI which correlated positively in both genotypes and again supports our finding that no major cognitive decline was observed in 5XFAD subjects. Finally, unique positive links were observed between HSP-related ER stress markers/CREB and total activity during OF only in controls, which were absent in the 5XFAD animals. Their relevance is not clear yet.

Collectively, our results point towards sustained hyperactivity during all phases of the test in 5X mice, this correlated well with levels of pathological and inflammatory markers in 5XFAD mice. Phenotypes were more pronounced in females, in line with observations in human patients. Conversely, neurotrophic factors and ER chaperones support cognitive performance and correlated with one another and were particularly prominent in male mice, possibly contributing to the alleviation of their phenotype.

## ACKNOWLEDGEMENTS

This project included funding from the Innovative Medicines Initiative 2/EFPIA, European Quality in Preclinical Data (EQIPD) consortium under grant agreement number 777364. The authors thank Dr. Heather Buchanan and Dr. Claire Hull for their valuable help with the molecular techniques and Jack Bray for his assistance with tissue collection. We would also like to acknowledge the staff of the Medical Research Facility for their support with animal care, handling and behavioural experiments and the qPCR Core Facility at the Institute of Medical Sciences for use of their qPCR systems.

## REFERENCES

[1] Prince M, Knapp M, Guerchet M, McCrone P, Prina M, Comas-Herrera A, Wittenberg R, Adelaja B, Hu B, King D, Rehill A, Salimkumar D (2014) Dementia UK: Second Edition - Overview, Alzheimer’s Society.

[2] Beam CR, Kaneshiro C, Jang JY, Reynolds CA, Pedersen NL, Gatz M (2018) Differences Between Women and Men in Incidence Rates of Dementia and Alzheimer’s Disease. J Alzheimers Dis 64, 1077–1083.

[3] Barnes LL, Wilson RS, Bienias JL, Schneider JA, Evans DA, Bennett DA (2005) Sex differences in the clinical manifestations of Alzheimer disease pathology. Arch Gen Psychiatry 62, 685–91.

[4] Koran MEI, Wagener M, Hohman TJ, Alzheimer’s Neuroimaging Initiative (2017) Sex differences in the association between AD biomarkers and cognitive decline. Brain Imaging Behav 11, 205– 213.

[5] Yan Y, Dominguez S, Fisher DW, Dong H (2018) Sex differences in chronic stress responses and Alzheimer’s disease. Neurobiol Stress 8, 120–126.

[6] Nichols E, Szoeke CEI, Vollset SE, Abbasi N, Abd-Allah F, Abdela J, Aichour MTE, Akinyemi RO, Alahdab F, Asgedom SW, Awasthi A, Barker-Collo SL, Baune BT, Béjot Y, Belachew AB, Bennett DA, Biadgo B, Bijani A, Bin Sayeed MS, Brayne C, Carpenter DO, Carvalho F, Catalá-López F, Cerin E, Choi J-YJ, Dang AK, Degefa MG, Djalalinia S, Dubey M, Duken EE, Edvardsson D, Endres M, Eskandarieh S, Faro A, Farzadfar F, Fereshtehnejad S-M, Fernandes E, Filip I, Fischer F, Gebre AK, Geremew D, Ghasemi-Kasman M, Gnedovskaya E V., Gupta R, Hachinski V, Hagos TB, Hamidi S, Hankey GJ, Haro JM, Hay SI, Irvani SSN, Jha RP, Jonas JB, Kalani R, Karch A, Kasaeian A, Khader YS, Khalil IA, Khan EA, Khanna T, Khoja TAM, Khubchandani J, Kisa A, Kissimova-Skarbek K, Kivimäki M, Koyanagi A, Krohn KJ, Logroscino G, Lorkowski S, Majdan M, Malekzadeh R, März W, Massano J, Mengistu G, Meretoja A, Mohammadi M, Mohammadi-Khanaposhtani M, Mokdad AH, Mondello S, Moradi G, Nagel G, Naghavi M, Naik G, Nguyen LH, Nguyen TH, Nirayo YL, Nixon MR, Ofori-Asenso R, Ogbo FA, Olagunju AT, Owolabi MO, Panda-Jonas S, Passos VM de A, Pereira DM, Pinilla-Monsalve GD, Piradov MA, Pond CD, Poustchi H, Qorbani M, Radfar A, Reiner RC, Robinson SR, Roshandel G, Rostami A, Russ TC, Sachdev PS, Safari H, Safiri S, Sahathevan R, Salimi Y, Satpathy M, Sawhney M, Saylan M, Sepanlou SG, Shafieesabet A, Shaikh MA, Sahraian MA, Shigematsu M, Shiri R, Shiue I, Silva JP, Smith M, Sobhani S, Stein DJ, Tabarés-Seisdedos R, Tovani-Palone MR, Tran BX, Tran TT, Tsegay AT, Ullah I, Venketasubramanian N, Vlassov V, Wang Y-P, Weiss J, Westerman R, Wijeratne T, Wyper GMA, Yano Y, Yimer EM, Yonemoto N, Yousefifard M, Zaidi Z, Zare Z, Vos T, Feigin VL, Murray CJL (2019) Global, regional, and national burden of Alzheimer’s disease and other dementias, 1990–2016: a systematic analysis for the Global Burden of Disease Study 2016. Lancet Neurol 18, 88–106.

[7] Snyder HM, Asthana S, Bain L, Brinton R, Craft S, Dubal DB, Espeland MA, Gatz M, Mielke MM, Raber J, Rapp PR, Yaffe K, Carrillo MC (2016) Sex biology contributions to vulnerability to Alzheimer’s disease: A think tank convened by the Women’s Alzheimer’s Research Initiative. Alzheimer’s Dement 12, 1186–1196.

[8] Yang J-T, Wang Z-J, Cai H-Y, Yuan L, Hu M-M, Wu M-N, Qi J-S (2018) Sex Differences in Neuropathology and Cognitive Behavior in APP/PS1/tau Triple-Transgenic Mouse Model of Alzheimer’s Disease. Neurosci Bull 34, 736–746.

[9] Carroll JC, Rosario ER, Kreimer S, Villamagna A, Gentzschein E, Stanczyk FZ, Pike CJ (2010) Sex differences in β-amyloid accumulation in 3xTg-AD mice: role of neonatal sex steroid hormone exposure. Brain Res 1366, 233–45.

[10] Sun Y, Guo Y, Feng X, Jia M, Ai N, Dong Y, Zheng Y, Fu L, Yu B, Zhang H, Wu J, Yu X, Wu H, Kong W (2020) The behavioural and neuropathologic sexual dimorphism and absence of MIP-3α in tau P301S mouse model of Alzheimer’s disease. J Neuroinflammation 17, 72.

[11] Plucińska K, Crouch B, Koss D, Robinson L, Siebrecht M, Riedel G, Platt B (2014) Knock-in of human BACE1 cleaves murine APP and reiterates alzheimer-like phenotypes. J Neurosci 34, 10710–10728.

[12] Platt B, Drever B, Koss D, Stoppelkamp S, Jyoti A, Plano A, Utan A, Merrick G, Ryan D, Melis V, Wan H, Mingarelli M, Porcu E, Scrocchi L, Welch A, Riedel G (2011) Abnormal Cognition, Sleep, EEG and Brain Metabolism in a Novel Knock-In Alzheimer Mouse, PLB1. PLoS One 6, e27068.

[13] Masters CL, Bateman R, Blennow K, Rowe CC, Sperling RA, Cummings JL (2015) Alzheimer’s disease. Nat Rev Dis Prim 1, 1–18.

[14] Selkoe DJ (2001) Alzheimer’s disease: Genes, proteins, and therapy. Physiol Rev 81, 741–766.

[15] Koss DJ, Platt B (2017) Alzheimer’s disease pathology and the unfolded protein response. Behav Pharmacol 28, 161–178.

[16] Cunningham C, Campion S, Lunnon K, Murray CL, Woods JFC, Deacon RMJ, Rawlins JNP, Perry VH (2009) Systemic Inflammation Induces Acute Behavioral and Cognitive Changes and Accelerates Neurodegenerative Disease. Biol Psychiatry 65, 304–312.

[17] Singhal G, Jaehne EJ, Corrigan F, Toben C, Baune BT (2014) Inflammasomes in neuroinflammation and changes in brain function: a focused review. Front Neurosci 8,.

[18] Dhawan G, Floden AM, Combs CK (2012) Amyloid-β oligomers stimulate microglia through a tyrosine kinase dependent mechanism. Neurobiol Aging 33, 2247–2261.

[19] Girard SD, Jacquet M, Baranger K, Migliorati M, Escoffier G, Bernard A, Khrestchatisky M, Féron F, Rivera S, Roman FS, Marchetti E (2014) Onset of hippocampus-dependent memory impairments in 5XFAD transgenic mouse model of Alzheimer’s disease. Hippocampus 24, 762–772.

[20] Ardestani PM, Evans AK, Yi B, Nguyen T, Coutellier L, Shamloo M (2017) Modulation of neuroinflammation and pathology in the 5XFAD mouse model of Alzheimer’s disease using a biased and selective beta-1 adrenergic receptor partial agonist. Neuropharmacology 116, 371– 386.

[21] Yang Y, Wang H, Kouadir M, Song H, Shi F (2019) Recent advances in the mechanisms of NLRP3 inflammasome activation and its inhibitors. Cell Death Dis 10, 128.

[22] Hull C, Dekeryte R, Buchanan H, Kamli-Salino S, Robertson A, Delibegovic M, Platt B (2020) NLRP3 inflammasome inhibition with MCC950 improves insulin sensitivity and inflammation in a mouse model of frontotemporal dementia. Neuropharmacology.

[23] Hetz C, Saxena S (2017) ER stress and the unfolded protein response in neurodegeneration. Nat Rev Neurol 13, 477–491.

[24] Liu B, Zhu Y, Zhou J, Wei Y, Long C, Chen M, Ling Y, Ge J, Zhuo Y (2014) Endoplasmic reticulum stress promotes amyloid-beta peptides production in RGC-5 cells. Cell Stress Chaperones 19, 827.

[25] Fonseca ACRG, Ferreiro E, Oliveira CR, Cardoso SM, Pereira CF (2013) Activation of the endoplasmic reticulum stress response by the amyloid-beta 1–40 peptide in brain endothelial cells. Biochim Biophys Acta - Mol Basis Dis 1832, 2191–2203.

[26] Pugazhenthi S, Wang M, Pham S, Sze C-I, Eckman CB (2011) Downregulation of CREB expression in Alzheimer’s brain and in Aβ-treated rat hippocampal neurons. Mol Neurodegener 6, 60.

[27] Zuccato C, Cattaneo E (2009) Brain-derived Neurotrophic Factor in Neurodegenerative Diseases. Nat Rev Neurol 5,.

[28] Oakley H, Cole SL, Logan S, Maus E, Shao P, Craft J, Guillozet-Bongaarts A, Ohno M, Disterhoft J, Van Eldik L, Berry R, Vassar R (2006) Intraneuronal β-amyloid aggregates, neurodegeneration, and neuron loss in transgenic mice with five familial Alzheimer’s disease mutations: Potential factors in amyloid plaque formation. J Neurosci 26, 10129–10140.

[29] Sadleir KR, Popovic J, Vassar R (2018) ER stress is not elevated in the 5XFAD mouse model of Alzheimer’s disease. J Biol Chem 293, 18434–18443.

[30] de Pins B, Cifuentes-Díaz C, Thamila Farah A, López-Molina L, Montalban E, Sancho-Balsells A, López A, Ginés S, Delgado-García JM, Alberch J, Gruart A, Girault J-A, Giralt A (2019) Conditional BDNF delivery from astrocytes rescues memory deficits, spine density and synaptic properties in the 5xFAD mouse model of Alzheimer disease. J Neurosci 39, 2121–18.

[31] Sadleir KR, Eimer WA, Cole SL, Vassar R (2015) Aβ reduction in BACE1 heterozygous null 5XFAD mice is associated with transgenic APP level. Mol Neurodegener 10,.

[32] Bundy JL, Vied C, Badger C, Nowakowski RS (2019) Sex-biased hippocampal pathology in the 5XFAD mouse model of Alzheimer’s disease: A multi-omic analysis. J Comp Neurol 527, 462–475.

[33] Creighton SD, Mendell AL, Palmer D, Kalisch BE, MacLusky NJ, Prado VF, Prado MAM, Winters BD (2019) Dissociable cognitive impairments in two strains of transgenic Alzheimer’s disease mice revealed by a battery of object-based tests. Sci Rep 9, 1–12.

[34] Landel V, Baranger K, Virard I, Loriod B, Khrestchatisky M, Rivera S, Benech P, Féron F (2014) Temporal gene profiling of the 5XFAD transgenic mouse model highlights the importance of microglial activation in Alzheimer’s disease. Mol Neurodegener 9, 33.

[35] O’Leary TP, Mantolino HM, Stover KR, Brown RE (2018) Age-related deterioration of motor function in male and female 5xFAD mice from 3 to 16 months of age. *Genes*, Brain Behav 1–11.

[36] O’Leary TP, Robertson A, Chipman PH, Rafuse VF, Brown RE (2018) Motor function deficits in the 12 month-old female 5xFAD mouse model of Alzheimer’s disease. Behav Brain Res 337, 256–263.

[37] Jawhar S, Trawicka A, Jenneckens C, Bayer TA, Wirths O (2012) Motor deficits, neuron loss, and reduced anxiety coinciding with axonal degeneration and intraneuronal Aβ aggregation in the 5XFAD mouse model of Alzheimer’s disease. Neurobiol Aging 33, 196.e29–196.e40.

[38] Flanigan TJ, Xue Y, Kishan Rao S, Dhanushkodi A, McDonald MP (2014) Abnormal vibrissa-related behavior and loss of barrel field inhibitory neurons in 5xFAD transgenics. Genes Brain Behav 13, 488–500.

[39] Kosel F, Torres Munoz P, Yang JR, Wong AA, Franklin TB (2019) Age-related changes in social behaviours in the 5xFAD mouse model of Alzheimer’s disease. Behav Brain Res 362, 160–172.

[40] Egan KJ, Vesterinen HM, Beglopoulos V, Sena ES, Macleod MR (2016) From a mouse: systematic analysis reveals limitations of experiments testing interventions in Alzheimer’s disease mouse models. Evidence-based Preclin Med.

[41] Flórez-Vargas O, Brass A, Karystianis G, Bramhall M, Stevens R, Cruickshank S, Nenadic G (2016) Bias in the reporting of sex and age in biomedical research on mouse models. Elife.

[42] Yeap J, Crouch B, Riedel G, Platt B (2020) Sequential habituation to space, object and stranger is differentially modulated by glutamatergic, cholinergic and dopaminergic transmission. Behav Pharmacol 31, 652–670.

[43] Percie du Sert N, Hurst V, Ahluwalia A, Alam S, Avey MT, Baker M, Browne WJ, Clark A, Cuthill IC, Dirnagl U, Emerson M, Garner P, Holgate ST, Howells DW, Karp NA, Lazic SE, Lidster K, MacCallum CJ, Macleod M, Pearl EJ, Petersen OH, Rawle F, Reynolds P, Rooney K, Sena ES, Silberberg SD, Steckler T, Würbel H (2020) The ARRIVE guidelines 2.0: Updated guidelines for reporting animal research. PLOS Biol 18, e3000410.

[44] Vollert J, Schenker E, Macleod M, Bespalov A, Wuerbel H, Michel M, Dirnagl U, Potschka H, Waldron A-M, Wever K, Steckler T, van de Casteele T, Altevogt B, Sil A, Rice ASC (2020) Systematic review of guidelines for internal validity in the design, conduct and analysis of preclinical biomedical experiments involving laboratory animals. BMJ Open Sci 4,.

[45] Koss DJ, Jones G, Cranston A, Gardner H, Kanaan NM, Platt B (2016) Soluble pre-fibrillar tau and β-amyloid species emerge in early human Alzheimer’s disease and track disease progression and cognitive decline. Acta Neuropathol 132, 875–895.

[46] Koss DJ, Dubini M, Buchanan H, Hull C, Platt B (2018) Distinctive temporal profiles of detergent-soluble and -insoluble tau and Aβ species in human Alzheimer’s disease. Brain Res 1699, 121– 134.

[47] Hull C, Dekeryte R, Koss DJ, Crouch B, Buchanan H, Delibegovic M, Platt B (2020) Knock-in of Mutated hTAU Causes Insulin Resistance, Inflammation and Proteostasis Disturbance in a Mouse Model of Frontotemporal Dementia. Mol Neurobiol 57, 539–550.

[48] Sampaio TB, Savall AS, Gutierrez MEZ, Pinton S (2017) Neurotrophic factors in Alzheimer’s and parkinson’s diseases: Implications for pathogenesis and therapy. Neural Regen Res.

[49] Chan CB, Ye K (2017) Sex differences in brain-derived neurotrophic factor signaling and functions. J Neurosci Res.

[50] Jiao S-S, Bu X-L, Liu Y-H, Zhu C, Wang Q-H, Shen L-L, Liu C-H, Wang Y-R, Yao X-Q, Wang Y-J (2016) Sex Dimorphism Profile of Alzheimer’s Disease-Type Pathologies in an APP/PS1 Mouse Model. Neurotox Res 29, 256–266.

[51] Laws KR, Irvine K, Gale TM (2016) Sex differences in cognitive impairment in Alzheimer’s disease. World J Psychiatry 6, 54.

[52] Paesler K, Xie K, Hettich MM, Siwek ME, Ryan DP, Schröder S, Papazoglou A, Broich K, Müller R, Trog A, Garthe A, Kempermann G, Weiergräber M, Ehninger D (2015) Limited effects of an eIF2α S51A allele on neurological impairments in the 5xFAD mouse model of Alzheimer’s disease. Neural Plast 2015,.

[53] Jyoti A, Plano A, Riedel G, Platt B (2010) EEG, activity, and sleep architecture in a transgenic AβPPswe/PSEN1A246E Alzheimer’s disease mouse. J Alzheimers Dis 22, 873–87.

[54] Robinson L, Riedel G (2014) Comparison of automated home-cage monitoring systems: Emphasis on feeding behaviour, activity and spatial learning following pharmacological interventions. J Neurosci Methods.

[55] Platt B, Riedel G (2011) The cholinergic system, EEG and sleep. Behav Brain Res 221,.

[56] Braun D, Feinstein DL (2019) The locus coeruleus neuroprotective drug vindeburnol normalizes behavior in the 5xFAD transgenic mouse model of Alzheimer’s disease. Brain Res 1702, 29–37.

[57] Kubota T, Matsumoto H, Kirino Y (2016) Ameliorative effect of membrane-associated estrogen receptor G protein coupled receptor 30 activation on object recognition memory in mouse models of Alzheimer’s disease. J Pharmacol Sci 131, 219–222.

[58] Howlett DR, Richardson JC, Austin A, Parsons AA, Bate ST, Davies DC, Gonzalez MI (2004) Cognitive correlates of Aβ deposition in male and female mice bearing amyloid precursor protein and presenilin-1 mutant transgenes. Brain Res 1017, 130–136.

[59] Hamm V, Héraud C, Bott JB, Herbeaux K, Strittmatter C, Mathis C, Goutagny R (2017) Differential contribution of APP metabolites to early cognitive deficits in a TgCRND8 mouse model of Alzheimer’s disease. Sci Adv 3,.

[60] Filali M, Lalonde R, Rivest S (2011) Anomalies in social behaviors and exploratory activities in an APPswe/PS1 mouse model of Alzheimer’s disease. Physiol Behav 104, 880–885.

[61] Maarouf CL, Kokjohn TA, Whiteside CM, Macias MP, Kalback WM, Sabbagh MN, Beach TG, Vassar R, Roher AE (2013) Molecular Differences and Similarities between Alzheimer’s Disease and the 5XFAD Transgenic Mouse Model of Amyloidosis. Biochem Insights 6, BCI.S13025.

[62] Heneka MT, Kummer MP, Stutz A, Delekate A, Schwartz S, Vieira-Saecker A, Griep A, Axt D, Remus A, Tzeng TC, Gelpi E, Halle A, Korte M, Latz E, Golenbock DT (2013) NLRP3 is activated in Alzheimer’s disease and contributes to pathology in APP/PS1 mice. Nature 493, 674–678.

[63] Halle A, Hornung V, Petzold GC, Stewart CR, Monks BG, Reinheckel T, Fitzgerald KA, Latz E, Moore KJ, Golenbock DT (2008) The NALP3 inflammasome is involved in the innate immune response to amyloid-β. Nat Immunol 9, 857–865.

[64] Tejera D, Mercan D, Sanchez-Caro JM, Hanan M, Greenberg D, Soreq H, Latz E, Golenbock D, Heneka MT (2019) Systemic inflammation impairs microglial Aβ clearance through NLRP 3 inflammasome. EMBO J 38,.

[65] Wu PJ, Hung YF, Liu HY, Hsueh YP (2017) Deletion of the Inflammasome Sensor Aim2 Mitigates Aβ Deposition and Microglial Activation but Increases Inflammatory Cytokine Expression in an Alzheimer Disease Mouse Model. Neuroimmunomodulation 24, 29–39.

[66] Buchanan H, Mackay M, Palmer K, Tothová K, Katsur M, Platt B, Koss DJ (2020) Synaptic Loss, ER Stress and Neuro-Inflammation Emerge Late in the Lateral Temporal Cortex and Associate with Progressive Tau Pathology in Alzheimer’s Disease. Mol Neurobiol 57, 3258–3272.

[67] Shao CY, Mirra SS, Sait HBR, Sacktor TC, Sigurdsson EM (2011) Postsynaptic degeneration as revealed by PSD-95 reduction occurs after advanced Aβ and tau pathology in transgenic mouse models of Alzheimer’s disease. Acta Neuropathol 122, 285–92.

[68] Kim J, Kim J, Huang Z, Goo N, Bae HJ, Jeong Y, Park HJ, Cai M, Cho K, Jung SY, Bae SK, Ryu JH (2019) Theracurmin ameliorates cognitive dysfunctions in 5XFAD mice by improving synaptic function and mitigating oxidative stress. Biomol Ther 27, 327–335.

[69] Maiti P, Bowers Z, Bourcier-Schultz A, Morse J, Dunbar GL (2021) Preservation of dendritic spine morphology and postsynaptic signaling markers after treatment with solid lipid curcumin particles in the 5xFAD mouse model of Alzheimer’s amyloidosis. Alzheimer’s Res Ther 13, 37.

[70] Zeng Y, Zhang J, Zhu Y, Zhang J, Shen H, Lu J, Pan X, Lin N, Dai X, Zhou M, Chen X (2015) Tripchlorolide improves cognitive deficits by reducing amyloid β and upregulating synapse-related proteins in a transgenic model of Alzheimer’s Disease. J Neurochem 133, 38–52.

[71] Griñán-Ferré C, Marsal-García L, Bellver-Sanchis A, Kondengaden SM, Turga RC, Vázquez S, Pallàs M (2019) Pharmacological inhibition of G9a/GLP restores cognition and reduces oxidative stress, neuroinflammation and ß-Amyloid plaques in an early-onset Alzheimer’s disease mouse model. Aging (Albany NY) 11, 11591–11608.

[72] Bundy JL, Vied C, Nowakowski RS (2017) Sex differences in the molecular signature of the developing mouse hippocampus. BMC Genomics 18, 237.

[73] Wong HR (1999) Heat shock proteins. Facts, thoughts, and dreams. A. De Maio. Shock 11:1–12, 1999. *Shock* **12**, 323–5.

[74] Fernandez-Funez P, Sanchez-Garcia J, de Mena L, Zhang Y, Levites Y, Khare S, Golde TE, Rincon-Limas DE (2016) Holdase activity of secreted Hsp70 masks amyloid-β42 neurotoxicity in Drosophila. Proc Natl Acad Sci U S A 113, E5212–21.

[75] Rivera I, Capone R, Cauvi DM, Arispe N, De Maio A (2018) Modulation of Alzheimer’s amyloid β peptide oligomerization and toxicity by extracellular Hsp70. Cell Stress Chaperones 23, 269–279.

[76] Lyon MS, Milligan C (2019) Extracellular heat shock proteins in neurodegenerative diseases: New perspectives. Neurosci Lett 711, 134462.

[77] Cui W, Wang S, Wang Z, Wang Z, Sun C, Zhang Y (2017) Inhibition of PTEN Attenuates Endoplasmic Reticulum Stress and Apoptosis via Activation of PI3K/AKT Pathway in Alzheimer’s Disease. Neurochem Res 42, 3052–3060.

[78] Reinhardt S, Schuck F, Grösgen S, Riemenschneider M, Hartmann T, Postina R, Grimm M, Endres K (2014) Unfolded protein response signaling by transcription factor XBP-1 regulates ADAM10 and is affected in Alzheimer’s disease. FASEB J 28, 978–997.

[79] Yang J-T, Wang Z-J, Cai H-Y, Yuan L, Hu M-M, Wu M-N, Qi J-S (2018) Sex Differences in Neuropathology and Cognitive Behavior in APP/PS1/tau Triple-Transgenic Mouse Model of Alzheimer’s Disease. Neurosci Bull 34, 736–746.

