## Supplementary material for "Sex differences in behaviour and molecular pathology in the 5XFAD model": Suplementary Information

### Supplementary Information

**SUPPLEMENTARY TABLE S1**

**COMPLETE RESULTS OF 3-WAY ANOVA FOR OF-NO-SI**

#### OPEN FIELD

##### ACTIVITY IN THE OPEN FIELD (DISTANCE TRAVELLED)

3-way ANOVA, with sex, genotype, trial (RM) as factors

| Factor | F value | P-value |
|---|---|---|
| Trial | F (1, 34) = 182.0 | P<0.0001 |
| Sex | F (1, 34) = 2.853 | P=0.1003 |
| Genotype | F (1, 34) = 18.93 | P=0.0001 |
| Trial x Sex | F (1, 34) = 3.431 | P=0.0727 |
| Trial x Genotype | F (1, 34) = 0.4296 | P=0.5166 |
| Sex x Genotype | F (1, 34) = 0.1267 | P=0.7241 |
| Trial x Sex x Genotype | F (1, 34) = 15.06 | P=0.0005 |

##### TIME IN WALL ZONE

3-way ANOVA, with sex, genotype, trial (RM) as factors

| Factor | F value | P-value |
|---|---|---|
| Trial | F (1, 34) = 46.92 | P<0.0001 |
| Sex | F (1, 34) = 5.406 | P=0.0262 |
| Genotype | F (1, 34) = 1.947 | P=0.1720 |
| Trial x Sex | F (1, 34) = 0.007340 | P=0.9322 |
| Trial x Genotype | F (1, 34) = 1.180 | P=0.2851 |
| Sex x Genotype | F (1, 34) = 6.190 | P=0.0179 |
| Trial x Sex x Genotype | F (1, 34) = 0.0006814 | P=0.9793 |

#### NOVEL OBJECT

##### NO: TIME IN ZONE

3-way ANOVA, with sex, genotype, trial (RM) as factors

| Factor | F value | P-value |
|---|---|---|
| Trial | F (1, 34) = 15.14 | P=0.0004 |
| Sex | F (1, 34) = 8.759 | P=0.0056 |
| Genotype | F (1, 34) = 0.5173 | P=0.4769 |
| Trial x Sex | F (1, 34) = 2.232 | P=0.1444 |
| Trial x Genotype | F (1, 34) = 0.8432 | P=0.3649 |
| Sex x Genotype | F (1, 34) = 3.980 | P=0.0541 |
| Trial x Sex x Genotype | F (1, 34) = 0.8274 | P=0.3694 |

##### NO: NUMBER OF VISITS

3-way ANOVA, with sex, genotype, trial (RM) as factors

| Factor | F value | P-value |
|---|---|---|
| Trial | F (1, 34) = 232.2 | P<0.0001 |
| Sex | F (1, 34) = 0.4673 | P=0.4989 |
| Genotype | F (1, 34) = 11.15 | P=0.0020 |
| Trial x Sex | F (1, 34) = 3.064 | P=0.0891 |
| Trial x Genotype | F (1, 34) = 2.042 | P=0.1621 |
| Sex x Genotype | F (1, 34) = 0.1969 | P=0.6601 |
| Trial x Sex x Genotype | F (1, 34) = 2.042 | P=0.1621 |

NO: OVERALL ACTIVITY

3-way ANOVA, with sex, genotype, trial (RM) as factors

| Factor | F value | P-value |
|---|---|---|
| Trial | F (1, 34) = 317.9 | P<0.0001 |
| Sex | F (1, 34) = 1.314 | P=0.2596 |
| Genotype | F (1, 34) = 8.004 | P=0.0078 |
| Trial x Sex | F (1, 34) = 3.085 | P=0.0880 |
| Trial x Genotype | F (1, 34) = 3.420 | P=0.0731 |
| Sex x Genotype | F (1, 34) = 1.748 | P=0.1949 |
| Trial x Sex x Genotype | F (1, 34) = 1.444 | P=0.2378 |

#### SOCIAL INTERACTION

SI: TIME IN ZONE

3-way ANOVA, with sex, genotype, trial (RM) as factors

| Factor | F value | P-value |
|---|---|---|
| Trial | F (1, 34) = 18.69 | P=0.0001 |
| Sex | F (1, 34) = 1.123 | P=0.2967 |
| Genotype | F (1, 34) = 7.386 | P=0.0103 |
| Trial x Sex | F (1, 34) = 0.1888 | P=0.6667 |
| Trial x Genotype | F (1, 34) = 0.4639 | P=0.5004 |
| Sex x Genotype | F (1, 34) = 0.2600 | P=0.6134 |
| Trial x Sex x Genotype | F (1, 34) = 2.323 | P=0.1367 |

SI: NO OF VISITS

3-way ANOVA, with sex, genotype, trial (RM) as factors

| Factor | F value | P-value |
|---|---|---|
| Trial | F (1, 34) = 1.180 | P=0.2849 |
| Sex | F (1, 34) = 0.4670 | P=0.4990 |
| Genotype | F (1, 34) = 6.299 | P=0.0170 |
| Trial x Sex | F (1, 34) = 0.4534 | P=0.5053 |
| Trial x Genotype | F (1, 34) = 1.682 | P=0.2034 |

| | | |
|---|---|---|
| Sex x Genotype | F (1, 34) = 8.979 | P=0.0051 |
| Trial x Sex x Genotype | F (1, 34) = 0.02226 | P=0.8823 |

SI: OVERALL ACTIVITY

3-way ANOVA, with sex, genotype, trial (RM) as factors

| Factor | F value | P-value |
|---|---|---|
| Trial | F (1, 34) = 30.06 | P<0.0001 |
| Sex | F (1, 34) = 3.393 | P=0.0742 |
| Genotype | F (1, 34) = 1.365 | P=0.2508 |
| Trial x Sex | F (1, 34) = 6.665 | P=0.0143 |
| Trial x Genotype | F (1, 34) = 1.772 | P=0.1920 |
| Sex x Genotype | F (1, 34) = 3.057 | P=0.0894 |
| Trial x Sex x Genotype | F (1, 34) = 0.2716 | P=0.6057 |

#### SUPPLEMENTARY FIGURE LEGENDS

**Figure S1: Locomotor activity in the open field trials (OF1 and OF2):** Time course of distance traversed (in meters, m) during OF1 and OF2 trials for female (♀) **(A)** and male (♂) 5XFAD vs WT mice (**B**)**.** All data are group means +/- SD, with statistical outcomes of 2-way ANOVA with trial and time as repeated measures for each sex/genotype indicated below. Significances are indicated as: ***=p <0.001, **=p<0.01, *=p<0.05.

**Figure S2: Overall activity measures during NO and SI:** Total distance traversed (in meters, m, per trial) during NO (**A**) and SI (**B**) trials for female and male 5XFAD vs WT mice. Tables below graphs indicate statistically significant outcomes from a 3-way ANOVA with sex, genotype and trial (repeated measures) as factors. Full statistical details can be found in Suppl. 1. All data are group means +/- SD, Significances are indicated as: ****=p <0.0001, ***= p<0.001, **=p<0.01, *=p<0.05.

**Fig. S3: Coomassie loading controls:** Example images of Coomassie total protein stained nitrocellulose membranes, relating to examples of ECL images of western blots provided in main manuscript. Densitometric measurements were taken from each lane for normalisation of ECL measures specific to target protein. Note that some membranes were used for the detection of more than one antigen

(PSD-95 and GFAP) whereas Ponceau was used for the normalisation of Iba-1 as Coomassie did not bind very well to the membrane.

**FIGURE S1**

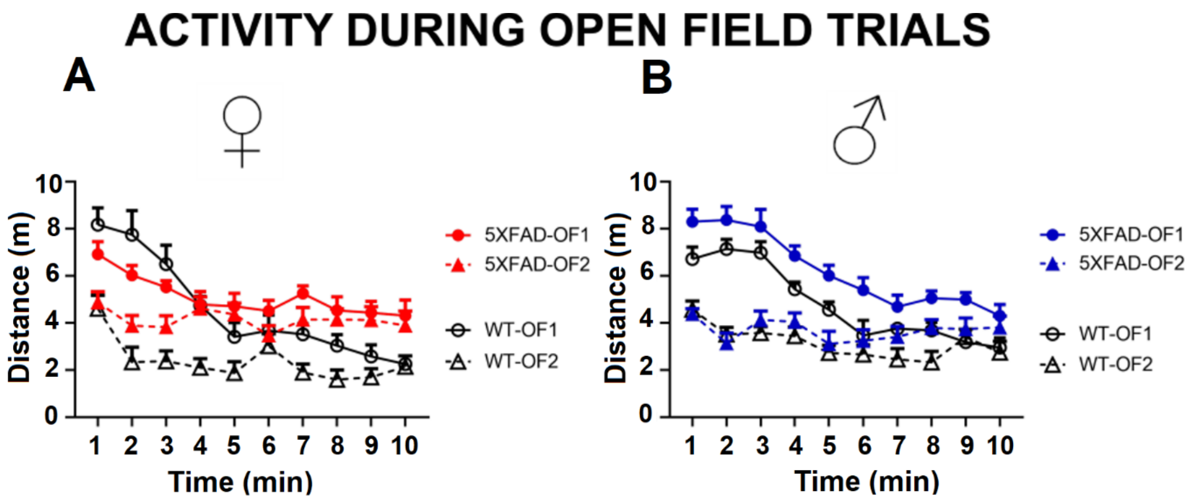


| Open Field | Within-trial habituation (OF1) | Between-trial habituation(OF2 vs OF1) |
|---|---|---|
| No habituation<br>Habituation | *Activity* | *Activity* |
| **Female WT** | *** | *** |
| **Male WT** | *** | *** |
| **Female Tg** | ** | * |
| **Male Tg** | *** | *** |

FIGURE S2

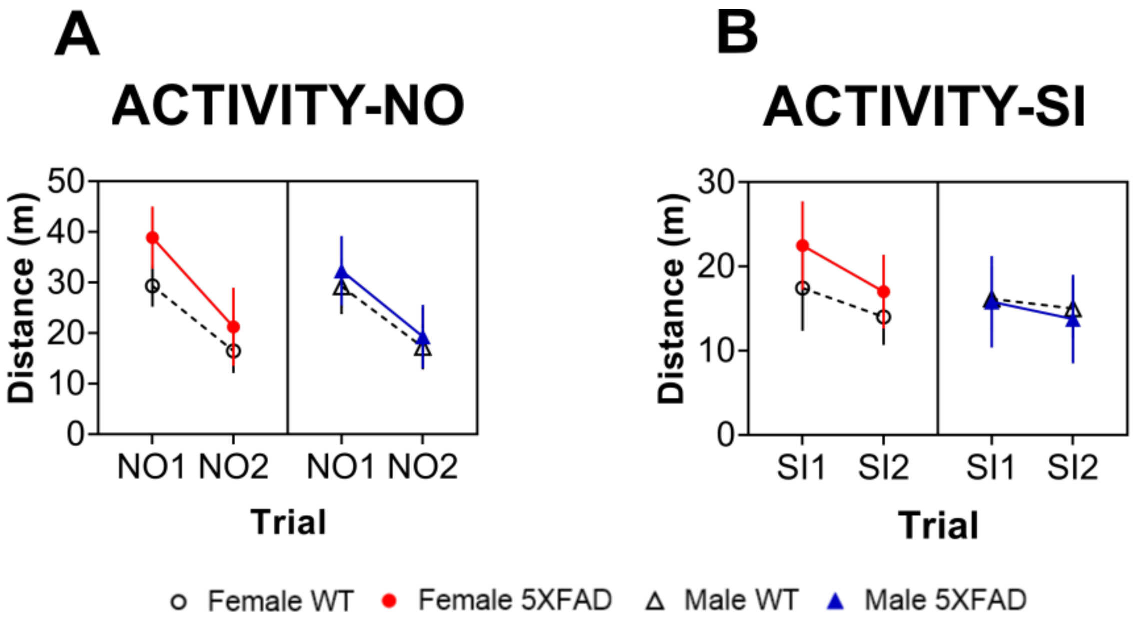


| | |
|---|---|
| **Trial** | **** |
| **Genotype** | ** |
| **Trial x sex** | *0.08* |
| **Trial x genotype** | *0.07* |

| | |
|---|---|
| **Trial** | *** |
| **Sex** | *0.07* |
| **Trial x sex** | * |
| **Sex x genotype** | *0.08* |

FIGURE S3

#### Coomassie total protein loading stain

6e10- soluble fraction

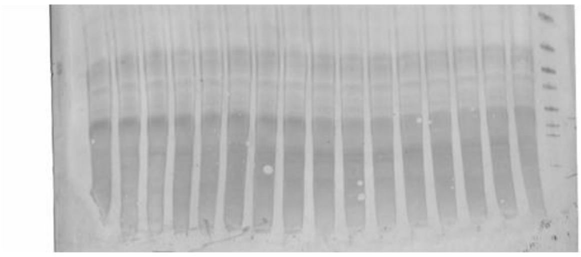

6e10- insoluble fraction

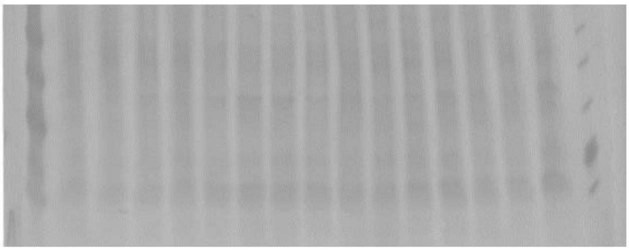

MOAB-2- soluble fraction

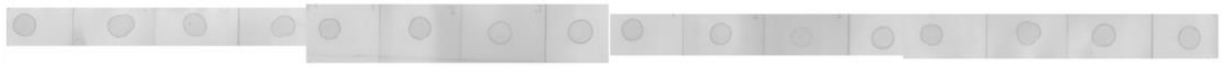

MOAB-2- insoluble fraction

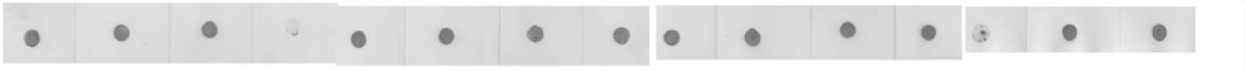

PSD-95

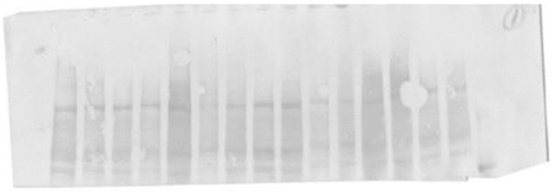

Synaptophysin

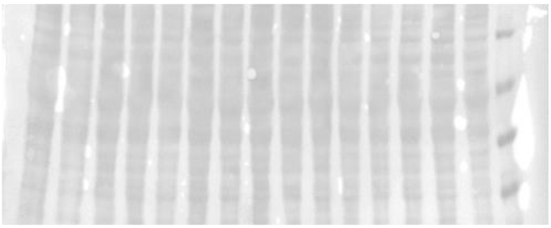

IBA1-Ponceau

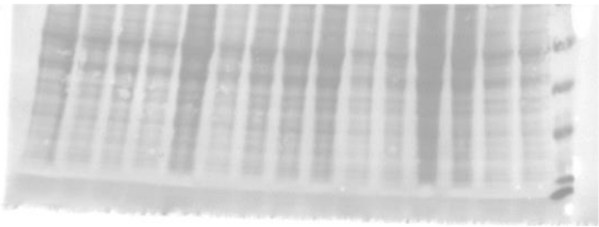

GFAP

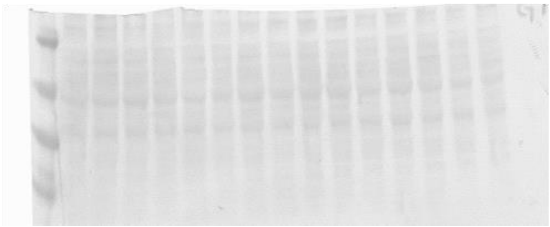
